# The Prioritization of Eleven-Nineteen-Leukemia Inhibitors as Potential Drug Candidates to Treat Acute Myeloid Leukemia

**DOI:** 10.1101/2022.12.07.519474

**Authors:** Xuejiao Shirley Guo, Peng-Hsun Chase Chen, Shiqing Xu, Wenshe Ray Liu

## Abstract

Acute myeloid leukemia (AML) is the second most diagnosed and the deadliest subtype of leukemia. Recently genetic loss-of-function studies have demonstrated that a human YEATS domain-containing protein named eleven-nineteen-leukaemia (ENL) functions as a transcriptional coactivator and is essential for the proliferation of AML that harbours oncogenic multiple lineage leukemia (MLL) rearrangements. We previously synthesized a series of small molecule inhibitors (**1**, **7-9**, **11-15** and **24**) that displayed significant and specific inhibitory effects against the ENL YEAST domain. In the current work, we report the development of a novel NanoBRET system that allows the analysis of cellular permeability, potency, selectivity, and stability of synthesized ENL inhibitors for their prioritization for further characterizations. Followed by *in vitro* metabolic stability and cell growth inhibition studies, we narrowed down to a potent and specific ENL YEATS domain inhibitor **13** with both high *in vitro* metabolic stability and strong anti-proliferation ability on MLL-fusion leukemia cell lines. A mouse pharmacokinetic (PK) analysis showed that at an oral dose of 20 mg/kg compound **13** had 60.9% bioavailability and 2.6 h mean residence time. With these favorable PK characteristics, compound **13** is ready for efficacy studies in an animal model. Cumulatively, the current study has prioritized compound **13** as a promising drug candidate to disrupt the pathogenic functions of ENL for the AML treatment.

## INTRODUCTION

Acute myeloid leukemia (AML) is the second most common type of leukemia diagnosed in adults and children and the deadliest subtype of leukemia in adults.^1^ Unlike chronic leukemia, acute leukemia progresses aggressively, and generally requires immediate treatment.^2^ However, the 5-year survival rate of patients with AML remains only around 25%, accounting for almost half of deaths from all subtypes of leukemia.^3^ AML is characterized by uncontrolled proliferation of abnormal myeloblasts which prevents the production of normal blood cells including mature red blood cells, neutrophils, monocytes, and platelets.^4–5^AML is usually associated with gene mutations, chromosomal rearrangements, and expressional alternations for multiple genes and microRNAs in myeloblasts.^6^ The majority of AML is typically associated with gene rearrangement caused by non-random chromosomal translocations.^7^ There are a total of 749 chromosomal aberrations in AML.^6^ Among these aberrations, the mixed-lineage leukemia gene (MLL or KMT2A) on the chromosomal 11q23 locus is one of the most frequent recurrent rearranged genes characterized in AML and is implicated in 5% ~ 10% of AML.^8^ The mutation in MLL usually associates with poor prognosis.^9^

By now, more than 80 genes have been reported to fuse with MLL, resulting in the production of chimeric proteins in which the *N*-terminal region of MLL is fused with the *C*-terminal part of a fusion protein partner.^10–13^ The most common five types of rearrangement in MLL-translocation-bearing AML are: MLL-AF4(t(4;11)(q21;q23)), MLL-AF9 (t(9;11)(p22;q23)), MLL-ENL (t(11;19)(q23;p13.3)), MLL-AF10 (t(10;11)(p12;q23)), and MLL-AF6(t(6;11)(q27;q23)).^14^ The AF9 and ENL fusion partner proteins share similar structure with an evolutionarily conserved YEATS protein family (named after the first-discovered members of this family: <u>Y</u>af9, <u>E</u>NL, <u>A</u>F9, <u>T</u>af14, and <u>S</u>as5).^15^ The YEATS domain works as an effective epigenetic reader for histone post-translational modifications including lysine acetylation and lysine crotonylation, leading to chromatin remodeling, transcriptional regulation and histone modification.^16–20^ The AF9 and ENL have been identified as key components of multiple transcriptional elongation complexes, such as super elongation complex (SEC) (Figure S1).^21^ In addition, leukemogenesis driven by MLL fusion proteins requires the histone methyltransferase DOT1L to trigger downstream gene transcriptions (e.g., the HOX/MEIS/MYC genes) for maintaining the self-reproductive state of transformed hematopoietic progenitor cells (Figure S1).^22–24^ Although AF9 and ENL are both subunits of the SEC/DOT1L transcriptional complex, recent studies identified ENL but not AF9 as a key factor to stabilize the association of SEC/Dot1L complex with DNA and maintaining the dysregulation of downstream genes, which is required for leukemogenesis.^26^ Notably, either disrupting the interaction between the ENL YEATS domain and acetylated histones or depleting the ENL protein could significantly suppress leukemia progression and has negligible effects on normal hematopoietic stem and progenitor cells as well.^25–26^ Moreover, the long and narrow hydrophobic “open-end” epigenetic reader pocket in the ENL YEATS domain (Figure 1) makes it a compelling target for drug discover for AML therapy, which was initially demonstrated in recent studies that developed both small compounds or peptides as AML inhibitors.^26^ For small molecule inhibitors, achieving high selectivity toward the ENL YEATS domain over the AF9 YEATS domain is challenging, as shown in several publications.^27–30^

**Figure 1.**
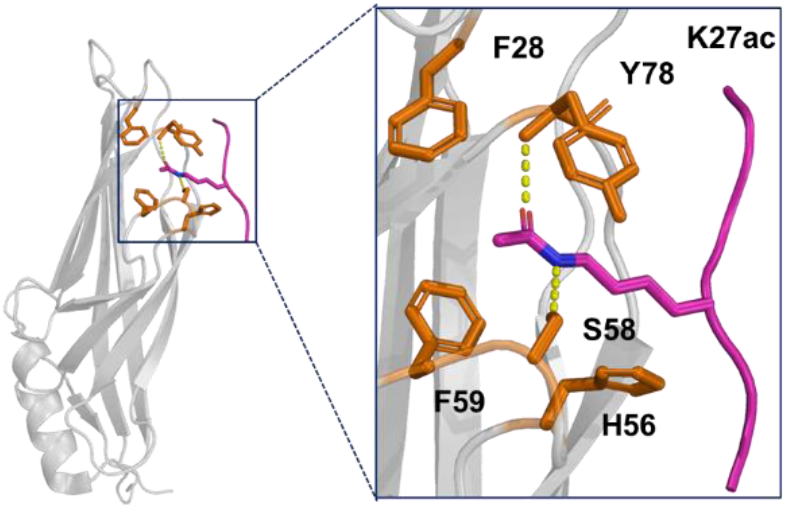
Overall structure of ENL YEATS domain bound to the H3K27ac. ENL YEATS is shown as gray ribbon with key residues of Kac pocket depicted as orange stick, acetyl-lysine in the H3K27ac ligand in the original crystal structure is colored in hot pink and its two hydrogen bonds with S58 and Y78 are colored in yellow.

In a previous study, we synthesized a series of potent small-molecule inhibitors that show preferential binding to the ENL YEATS domain rather than the AF9 YEATS domain *in vitro*.^31^ However, these inhibitors showed variable cellular potency in two MLL-rearranged leukemia cells, MOLM-13 and MV4-11, which is likely due to their differences in cellular permeability, stability, *etc*.^31^ To prioritize some of these inhibitors as potential drug leads for the treatment of AML and further optimization, in this study, we developed a fluorescence tracer based on a reported selective ENL/AF9 YEATS domain inhibitor SR-0813^32^ and constructed a NanoBRET system for the evaluation of cellular permeability, potency, selectivity, and stability of our developed ENL inhibitors. Potent inhibitors were then selected according to the IC_50_ values determined by the NanoBRET assay and subjected to *in vitro* metabolic stability, cell viability and anti-proliferation evaluation to screen inhibitors with optimal characteristics. One inhibitor, compound **13** was further advanced to the pharmacokinetic (PK) characterization in mice showing both high bioavailability and long mean residence time. We believe that the current study paves the foundation of exploring compound **13** in animal efficacy studies for treating AML and streamlines an applicable route for the exploration of next-generation AML inhibitors.

## RESULTS AND DISCUSSION

### Design, preparation, and validation of a NanoBRET system for the analysis of ENL YEATS domain inhibitors

Bioluminescent resonance energy transfer (BRET) refers to the non-radiative energy transfer from a luciferase energy donor to a fluorophore acceptor after the luciferase substrate oxidation, which only happens when there is very close proximity (<10 nm) between the donor and acceptor.^33–34^ It is a biophysical technique to monitor proximity of two components within live cells and has been used to investigate a range of biological processes including ligand binding, intracellular signaling, receptor-receptor proximity, and receptor transport.^35^ Over the years, researchers have used several luciferases for the BRET assay development including Firefly luciferase (Fluc), Renilla luciferase (Rluc), and an Rluc derivative Rluc8.^36–38^ A latest development is nanoluciferase (NLuc) that has a favorable small size with only a 19 kDa molecular weight. The development of NLuc and its substrate furimazine has significantly broadened applications of BRET assays due to the small luciferase size, good stability and high luminescence, resulting in the establishment of NanoBRET.^39^ Furthermore, recent studies suggested the possible use of NanoBRET to study the direct interaction of small molecules with intracellular targets.^40^ Apart from furimazine, the NanoBRET assay includes another two key components: an expressed cellular target protein fused with NLuc and a cellularly permeable fluorescent tracer that binds the target protein specifically. We set to design an ENL-NanoBRET system. In our ENL-NanoBRET design, the ENL YEATS domain was genetically fused at the *C*-terminus of NLuc and the fusion protein was recombinantly expressed in HEK293T cells. Since the ENL YEATS domain is also a small protein with a 16 kDa molecular weight, the two active sites in the ENL YEATS domain and NLuc are in a close distance. When a fluorescent tracer binds to the ENL YEATS domain, it will be in proximity to furimazine that binds to NLuc and consequently have a strong BRET signal. The BRET signal will be lost once a test compound binds to the ENL YEATS protein and displaces the tracer (Figure 2a). Since it was reported that the transiently expressed NanoBRET constructs showed substantial batch-to-batch variation,^41^ we generated HEK293T cell lines stably expressing NLuc-ENL YEATS to conduct the ENL-NanoBRET assays for better repeatability (Figure S2).

**Figure 2.**
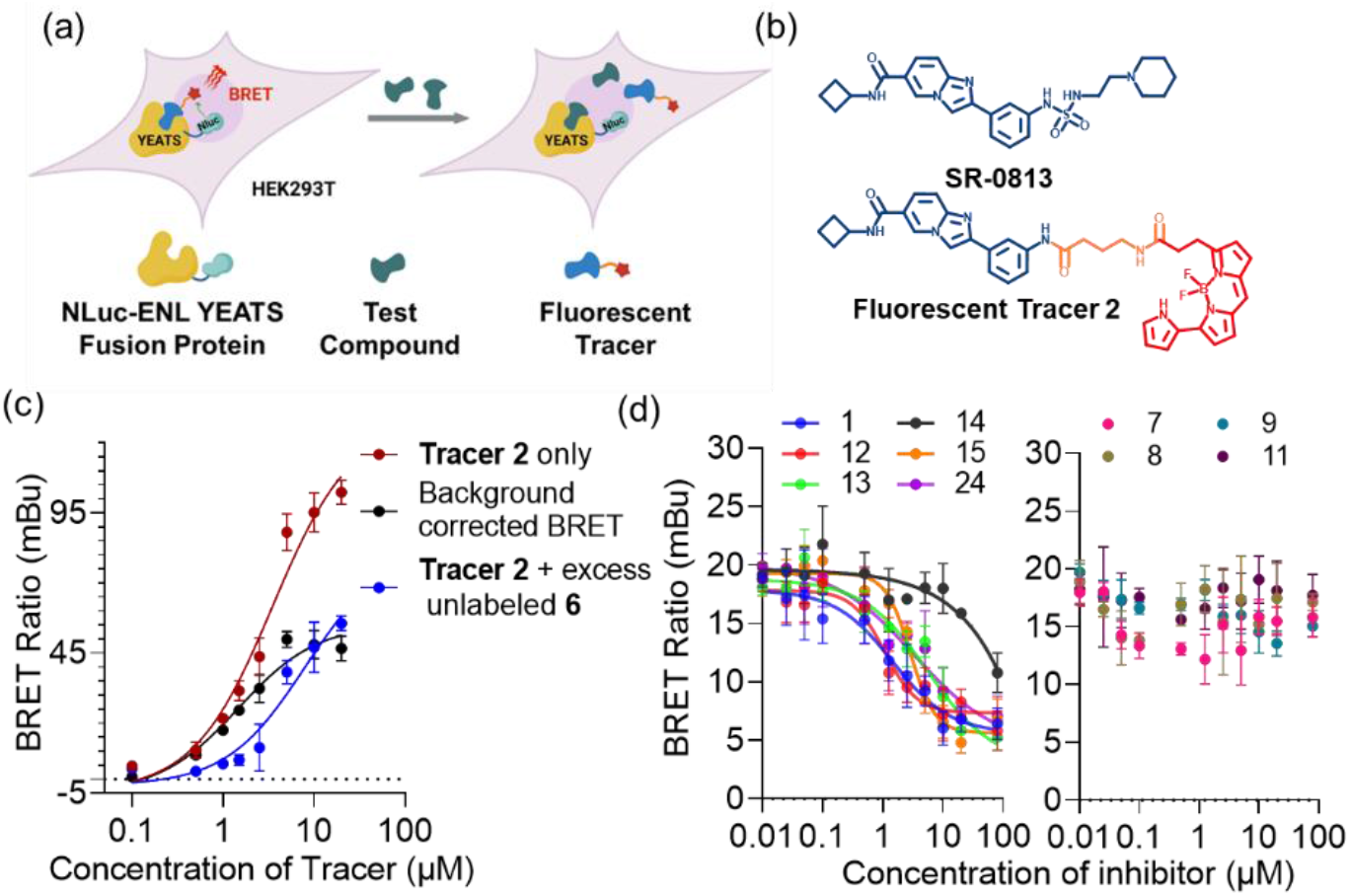
(a) Illustration of intracellular target engagement assay. A cell permeable fluorescent tracer binds in dynamic equilibrium to an intracellular target protein fused to NLuc, resulting in BRET. Introduction of compounds that bind the same target cause the tracer to be displaced, resulting in a decrease in BRET. (b) The ENL **Tracer 2** was derived from a ENL inhibitor (SR-0813, shown in blue) and the BODIPY590 dye (shown in red). (c) Apparent **Tracer 2** affinity for NLuc-ENL YEATS fusion protein in HEK293T cells. (d) NanoBRET curves of ENL inhibitor affinities for NLuc-ENL YEATS in HEK293T cells.

We firstly designed a NanoBRET **Tracer 1** based on inhibitor **1** (Scheme S1) by connecting a commonly used dye BODIPY590 through a butyl linker. Intracellular binding of this tracer to ENL YEATS domain was detected by measuring the bioluminescence emission from live cells. However, the profile of apparent affinity of **Tracer 1** for the NLuc-ENL YEATS fusion protein (Figure S3) limiting its application for further NanoBRET assays due to the low background corrected BRET values. Garnar-Wortzel *et al*.^32^ discovered a potent and selective ENL/AF9 YEATS domain inhibitor, SR-0813, (IC_50_ = 25 nM), which capable of disrupting the pathogenic function of ENL in acute leukemia. However, the rapid metabolism of SR-0813 in mouse liver microsomes [half-life time (t_1/2_) = 9.3 min] restricts its potential as an *in vivo* therapeutic.^42^ Inspired by this highly potent ENL inhibitor SR-0813 (Figure 2b, Top), we developed another cell-permeable small-molecule fluorescent **Tracer 2** (Figure 2b, Bottom) as a NanoBRET acceptor by liking BODIPY590 (Figure 2b, Bottom in red) to the core skeleton of SR-0813 (Figure 2b, Bottom in blue) with a butyl chain (Figure 2b, Bottom in orange) through two sequential amidation reactions. The following characterization showed successfully a saturable curve clearly dependent on the concentration of the **Tracer 2** (Figure 2c). The non-specific binding was eliminated by co-incubation with 80 μM of the unlabeled compound **6** (Scheme S2) to obtain the background-corrected BRET data (Figure 2c). Half-maximal stimulation (EC_50_) was observed at 1.46 μM. Similar to other competitive binding assays, the concentration of fluorescence tracer in the NanoBRET assay affects the apparent affinity of competitive compounds towards the ENL YEATS domain since both the tracer and test compounds competitively bind with the NLuc-ENL YEATS protein. A high concentration of tracer will increase the apparent IC_50_ value of tested inhibitors, which termed as the Cheng-Prusoff relationship.^43^ To optimize the fluorescence tracer concentration for the ENL-NanoBRET assay, the IC_50_ value of inhibitor **24** towards the ENL YEATS domain was evaluated in a competitive ENL-NanoBRET assay using various concentrations of **Tracer 2** (Figure S4). The quality of this competitive ENL inhibitor displacement assay was determined using the raw fold change in the BRET ratio observed at the 1 μM of **Tracer 2** compared to the BRET ratio in the presence of a saturating dose of unlabeled compound **6**, that defined the assay window (value = 3.59), together with the high Z factor (value = 0.62). A final **Tracer 2** concentration at 1 μM was determined as optimal for the ENL-NanoBRET assay.

With the optimal assay conditions in hand, we performed the competitive ENL-NanoBRET assays for the selected ENL inhibitors including **1**, **7**-**9**, **11**-**15** and **24** to evaluate their cellular ENL YEATS inhibition potency by co-incubating gradient concentrations of these inhibitors and 1 μM of tracer with HEK293T cells expressing NLuc-ENL YEATS for 2 h. Among these compounds, inhibitors **1** and **12** showed the most significantly BRET signal decrements with IC_50_ values as 1.42 μM and 1.13 μM respectively, indicating their high potency in binding the ENL YEATS active site in the cellular environment (Figure 2d, Left). Inhibitors **13**, **15** as well as **24** showed a comparable slightly lower inhibition potency toward ENL YEATS with IC_50_ values as 5.11 μM, 2.80 μM and 3.37 μM, respectively, which are in line with our previous *in vitro* biochemical affinity results. Compared to inhibitor **13**, its analogue **14** showed a relatively lower cellular inhibition potency toward ENL YEATS (Figure 2d, Left, IC_50_ value higher than 20 μM), although it binds the His-ENL *in vitro* with a similar potency as **13** (*in vitro* AlphaScreen IC_50_ values of **13** and **14** are 266 nM and 264 nM, respectively^31^). In addition, while inhibitors **7**-**9** and **11** displayed high potency determined by an *in vitro* AlphaScreen assay, these inhibitors were substantially less potent than the other inhibitors determined by the competitive ENL-NanoBRET assay. As shown in the right panel of Figure 2d, inhibitors **7**-**9** and **11** showed limited NanoBRET signal reductions at concentration up to 80 μM, indicating the weak cellular permeability or active efflux of these inhibitors. Our results so far prioritized 5 potent inhibitors (**1**, **12, 13**, **15**, and **24**) with high cellular potency for further characterizations.

### Metabolic stability of inhibitors 1, 12, 13, 14, 15, and 24 in human plasma and liver microsome

To test the potential *in vivo* applications for our prioritized inhibitors, we next moved to study their *in vitro* metabolic properties. As one of *in vitro* ADME (Absorption, Distribution, Metabolism, and Excretion) screening studies, the plasma stability assay is used to determine the stability of potential drugs in plasma.^44^ In general, the excellent *in vivo* efficacy depends on the slow degradation of the drug in plasma.^45^ More importantly, the drug stability assay in plasma could provide more accurate *in vitro* assessment of compounds. For those compounds containing esters, amides, lactones, lactams, carbamides or sulfonamides functional group, their plasma stability must be taken into consideration, since these groups are highly susceptible to hydrolysis in plasma.^46–47^ As shown in Figure 3a, stability in human plasma was tested for all inhibitors, **13**, **14** and **15** shown no significant degradation when incubated in human plasma at 37 °C for 2 h. These three compounds also showed ~95% remaining in plasma, suggesting the applaudable stabilities of them (Table 2). In contrast, inhibitors **1**, **12** and **24** showed slightly lower human plasma stability, with ~15% degradation after 2 h incubation (Figure 3a and Table 2).

**Figure 3.**
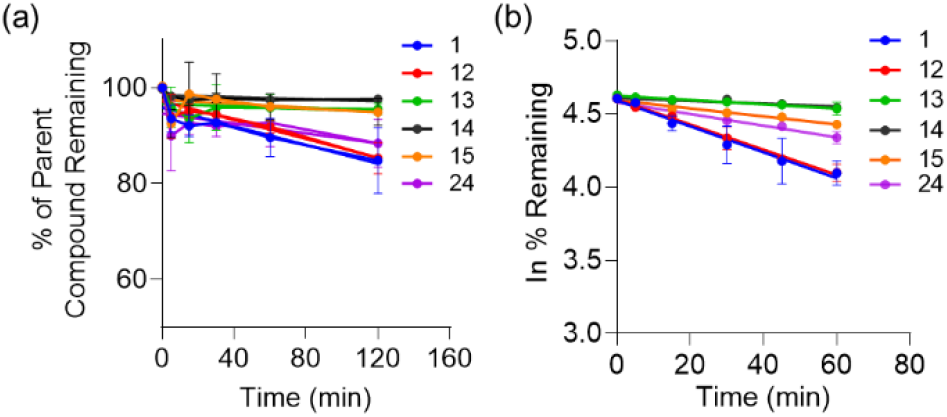
(a) Stability of ENL inhibitors in human plasma and human liver microsomes.

**Table 1.**
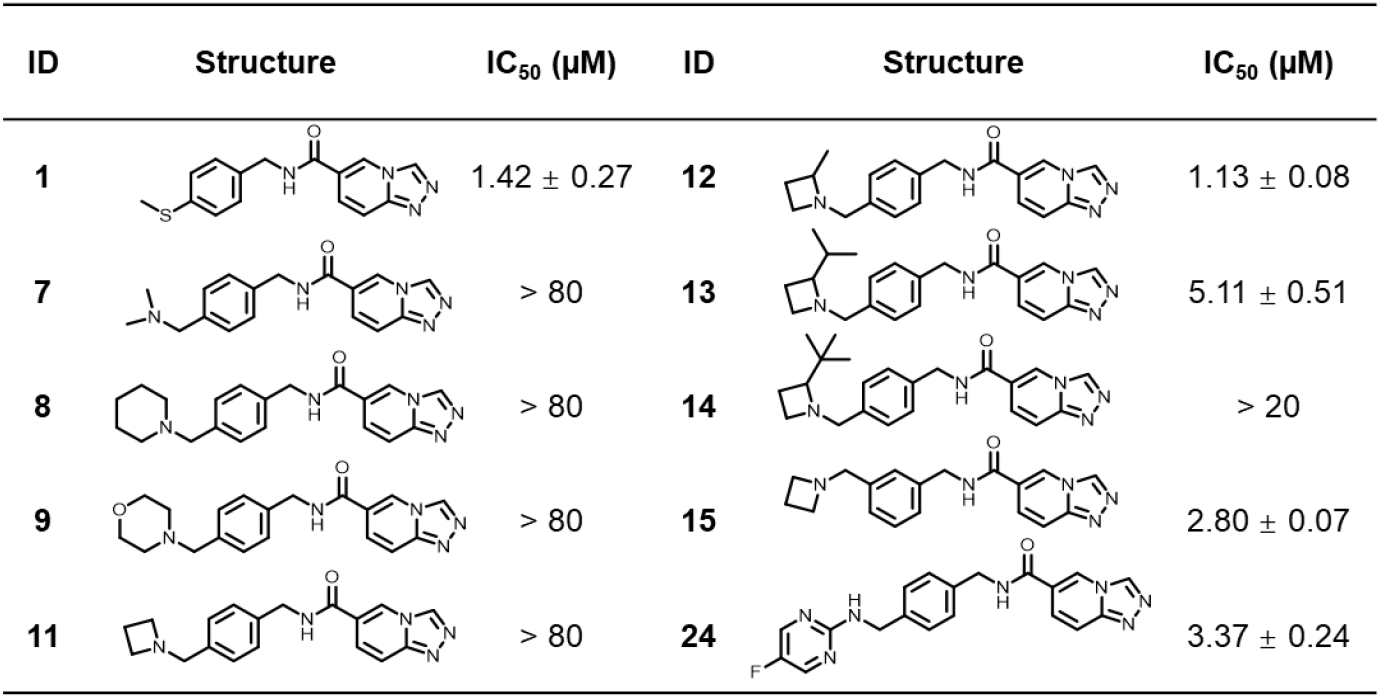
Cellular IC_50_ Values of ENL inhibitors Determined by NanoBRET.

**Table 2.** Summarized metabolic stability values for ENL inhibitors.

| ID | Plasma | Liver Microsome |  |
| --- | --- | --- | --- |
| | % of parent compound remaining (after 2h incubation) | $t_{1/2}$ (min) | CL<sub>int</sub>, pred (mL/min/kg) |
| **1** | 84.81 ± 6.84 | 80.65 ± 17.50 | 22.23 ± 4.76 |
| **12** | 85.32 ± 3.24 | 81.01 ± 7.02 | 21.56 ± 1.89 |
| **13** | 95.58 ± 1.20 | 426.80 ± 39.81 | 4.09 ± 0.38 |
| **14** | 97.60 ± 0.74 | 718.81 ± 44.47 | 2.42 ± 0.14 |
| **15** | 94.94 ± 2.81 | 254.05 ± 36.17 | 6.93 ± 1.01 |
| **24** | 88.42 ± 4.98 | 175.42 ± 13.22 | 9.95 ± 0.77 |

Hepatic metabolism is one of the major routes for drug elimination. *In vitro* hepatic metabolism studies are typically conducted using the microsomal incubation to determine the intrinsic clearance (CL_int_) rate.^48^ This assay is important for the drug development to screen leads from numerous compounds showed similar therapeutic potentials. To illustrate this point, human liver microsome including multiple important drug metabolizing enzymes, such as flavin monooxygenases (FMO), glucuronosyltransferases and esterase, were used for *in vitro* hepatic clearance determination among selected ENL inhibitors.^49^ The individual inhibitor was incubated with liver microsomes in either presence or absence of nicotinamide adenine dinucleotide phosphate (NADPH, the cofactor for FMO-dependent oxidations) at 37 °C. The degradations of inhibitors were detected via the LC-MS/MS analysis. As shown in Figure 3b, the metabolic stability of each inhibitor was determined by plotting natural logarithm (Ln) inhibitor remaining (y-axis) versus the incubation time (x-axis), where the corresponding slope represented the inhibitor metabolism rate. The slope values were then applied to calculate the in vitro t_1/2_ and intrinsic clearance rates that are presented in Table 2.

While inhibitors **1** and **12** degraded quickly in the microsomal stability assay with only ~ 60% remaining after 2 h incubation, inhibitor **24** showed a moderate degradation with acceptable intrinsic clearance rate (CLint = 9.95 mL/min/kg) (Table 2). In contrast, inhibitors **13**, **14**, and **15** all showed favorable microsomal stabilities with > 90% parent inhibitor remaining after 2 h incubation (4.09, 2.42 and 6.93 mL/min/kg respectively), suggested low intrinsic clearance rates for these three inhibitors (Table 2).

### The ENL is required for growth of acute leukaemia cell

After testing the *in vitro* metabolic stabilities of inhibitors, we next focused the characterization of their effects in killing tumor cells. Two cell lines (MV4-11 and MOLM-13) that are sensitive to the loss of ENL and the Jurkat cell line that is insensitive to the loss of ENL were used in the cellular evaluation.^25^ HEK293T cells were used as a non-tumor control as well.

We started with testing the viability of those four cell lines after 72 h inhibitor treatments. While slight cytotoxicity towards MOLM-13 (Figure 4a) and MV4-11(Figure 4b) cells was observed for inhibitors **1**, **12**, **14**, **15**, and **24**, inhibitor **13** showed the most significant cytotoxicity with determined IC_50_ values as 8.20 μM for MOLM-13 cells and 9.15 μM for MV4-11 cells. As shown in Figures 4c and 4d, the cytotoxicity of all inhibitors towards Jurkat or HEK293T cells are close to negligible at as high as 10 μM concentration. The distinct cell death was not observed until the inhibitor **13** concentration reached to 16 μM (Figure 4e).

**Figure 4.**
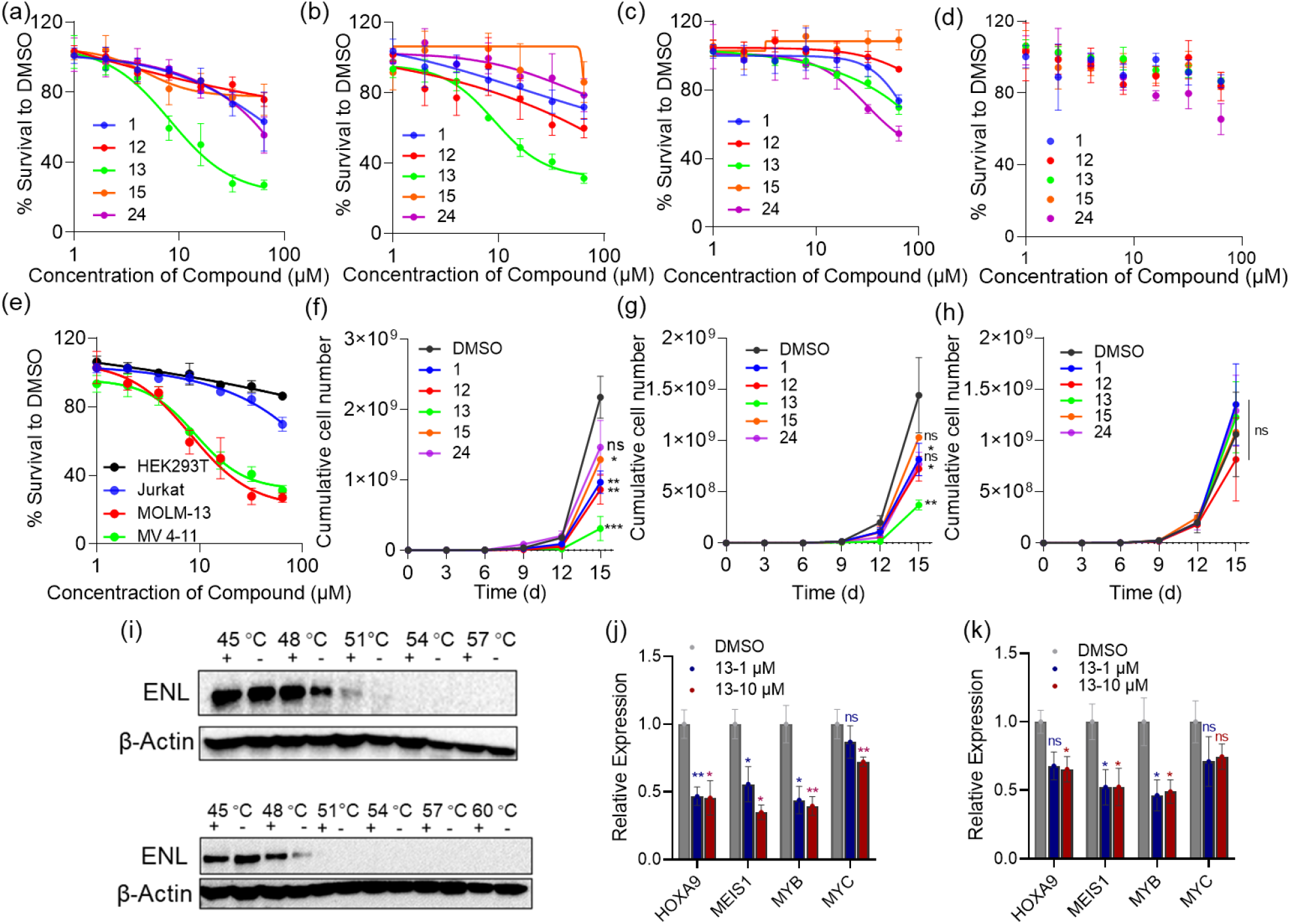
(a) MOLM-13 (b) MV4-11 (c) Jurkat and (d) HEK293T cell viability after 72-h ENL inhibitors treatment. (e) Comparison of cell viability among MOLM-13, MV4-11, Jurkat and HEK293T cell after 72-h inhibitor **13** treatment. Proliferation of (f) MOLM-13 (g) MV4-11 and (h) Jurkat cell in response to ENL inhibitor in 10 μM. (i) CETSAs in MOLM-13 (Top) and MV4-11 (Bottom) cells treated with **13** in 10 μM (+) or DMSO (-) at the indicated temperatures. β-Actin was used as a loading control. qRT-PCR analysis of HOXA9, MEIS1, MYB and MYC gene expression in (j) MOLM-13 and (k) MV4-11 cells treated with **13** or the DMSO negative control. *P < 0.05, **P < 0.01, ***P < 0.001, ****P < 0.0001. Not significant (n.s.) P > 0.05.

Based on the previous studies, ENL functions as a transcriptional activator to dysregulate gene expression, therefore supports the pathogenesis of acute leukemia. ^25^ Therefore, longer incubation time for ENL inhibitors may be essential for showing their antileukemic effects. To validate this prospect, the cell proliferation in the presence of ENL inhibitors was recorded for ~2 weeks to examine the anti-proliferation effect of them with extended treatments. Consistent with the cytotoxicity assays, inhibitor **13** remarkably inhibited the growth of two ENL-dependent cell lines, MOLM-13 (Figure 4f) and MV4-11(Figure 4g) at the concentration of 10 μM. It was also observed that inhibitor **13** was capable of inhibiting MOLM-13 and MV4-11 cell proliferation at low concentration as 1 μM (Figure S5a and S5b). In contrast, the co-incubation of **13** has no effect on the growth of Jurkat cell (Figure 4h and S5c). Moreover, cell cycle analysis data showed the increased G1 stage for MOLM-13 and MV4-11 cells after their treatment with inhibitor **13** (Figure S6), which is in line with the notion that loss of ENL results in G1 arrest during cell division.^26^

To confirm whether the growth inhibition effect was caused by on-target inhibition of the endogenous ENL protein, cellular thermal shift assay (CETSA) was carried to evaluate thermal stability of the ENL protein in MOLM-13 cells MV4-11 cells after the treatment with 10 μM of inhibitor **13** for 3 h and 6 h. In both cell lines, higher abundance of ENL protein was observed in **13**-treated than DMSO-treated cells at increased temperatures, suggesting the specific engagement of **13** with ENL within live cells (Figure 4i). Meanwhile the expression of ENL target genes including HOXA9, MEIS1, MYB and MYC in MOLM-13 and MV4-11 cells was evaluated after co-incubation with inhibitor **13** for 72 h. It was observed that inhibitor **13** effectively suppressed HOXA9, MEIS1 and MYB gene expression at as low as the concentration of 1.0 μM (Figure 4j and 4k).

Collectively, these data in accordance with the cell sensitivity caused by genetic disruption of ENL supports the idea that on-target effects are responsible for inhibitor **13**-mediated ENL-dependent leukemia growth inhibition.

### Pharmacokinetics Study

Inhibitor **13** was further advanced to a pharmacokinetic (PK) study in CD-1 mice that was carried out in Wuxi AppTec Inc. A dose of 20 mg/kg was conducted for both intravenous (IV) and oral (PO) administration. Inhibitor **13** exhibited t_1/2_ as 1.05 and 0.84 h, respectively, for IV and PO adminstrations. Its C_max_ values for IV and PO adminstrations were 12,058 and 2,080 ng/ml, resepectively. The oral C_max_ corresponds to 5.73 μM, an effective concentration to show as antiproliferation effect. What are remarkable for inhibitor **13** are its high bioavailability as 60.9% and relatively long mean residence time as 2.6 h. The high bioavailability and long residence time provide the opportunity to deliver high and sustained concentrations of the drug through oral administration (Figure 5). These favorable PK features pave the way for the anti-leukemia animal efficacy study of inhibitor **13**.

**Figure 5.**
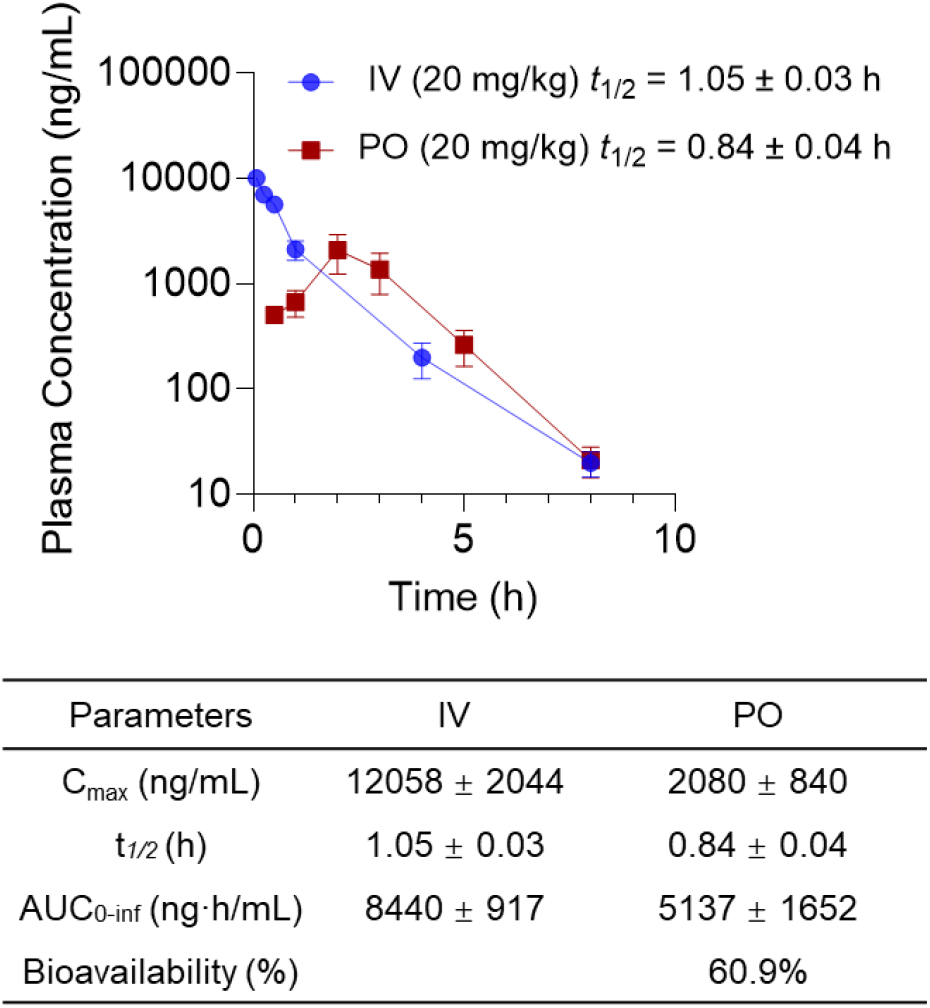
Plasma concentration of inhibitor **13** (Top) following IV or PO administration at dose of 20 mg/kg and corresponding characterization values (Bottom)(n=3).

## CONCLUSIONS

In summary, we have successfully developed an ENL-NanoBRET assay system for the cellular potency analysis of ENL inhibitors and used this novel assay system to prioritize five inhibitors **1**, **12**, **13**, **15**, and **24** for further characterizations. By examining metabolic stability in human plasma and liver microsome, we found that inhibitor **13** displayed lower elimination rate and better bioavailability compared with other inhibitors. More importantly, inhibitor **13** exhibited on-target effect in inhibiting leukemia cell growth. An *in vivo* PK analysis showed a high oral bioavailability and relative long mean residence time for inhibitor **13**. Collectively, our study prioritized inhibitor **13** as a prospective drug candidate to disrupt the pathogenic functions of ENL for AML treatment and a lead compound for further structure-activity relationship optimization.

## ACKNOWLEDGEMENTS

The project was supported in part by National Institutes of Health (Grants NIH-R21CA267512 and R35GM145351) and Welch Foundation (Grant A-1715). X. Guo was supported by a postdoctoral fellowship from Cancer Prevention and Research Institute of Texas (Grant RP210043).

## REFERENCE

1. De Kouchkovsky, I.; Abdul-Hay, M., Acute myeloid leukemia: a comprehensive review and 2016 update. Blood cancer journal 2016, 6 (7), e441–e441.

2. Ferrara, F.; Schiffer, C. A., Acute myeloid leukaemia in adults. The Lancet 2013, 381 (9865), 484–495.

3. Döhner, H.; Estey, E.; Grimwade, D.; Amadori, S.; Appelbaum, F. R.; Büchner, T.; Dombret, H.; Ebert, B. L.; Fenaux, P.; Larson, R. A., Diagnosis and management of AML in adults: 2017 ELN recommendations from an international expert panel. Blood, The Journal of the American Society of Hematology 2017, 129 (4), 424–447.

4. Tallman, M. S.; Gilliland, D. G.; Rowe, J. M., Drug therapy for acute myeloid leukemia. Blood 2005, 106 (4), 1154–1163.

5. Burnett, A.; Wetzler, M.; Lowenberg, B., Therapeutic advances in acute myeloid leukemia. J Clin Oncol 2011, 29 (5), 487–494.

6. Kumar, C. C., Genetic abnormalities and challenges in the treatment of acute myeloid leukemia. Genes & cancer 2011, 2 (2), 95–107.

7. Döhner, K.; Döhner, H., Molecular characterization of acute myeloid leukemia. Haematologica 2008, 93 (7), 976–982.

8. Shih, L.; Liang, D.; Fu, J.; Wu, J.; Wang, P.; Lin, T.; Dunn, P.; Kuo, M.; Tang, T.; Lin, T., Characterization of fusion partner genes in 114 patients with de novo acute myeloid leukemia and MLL rearrangement. Leukemia 2006, 20 (2), 218–223.

9. Patel, J. P.; Gönen, M.; Figueroa, M. E.; Fernandez, H.; Sun, Z.; Racevskis, J.; Van Vlierberghe, P.; Dolgalev, I.; Thomas, S.; Aminova, O., Prognostic relevance of integrated genetic profiling in acute myeloid leukemia. New England Journal of Medicine 2012, 366 (12), 1079–1089.

10. Munoz, L.; Nomdedeu, J.; Villamor, N.; Guardia, R.; Colomer, D.; Ribera, J.; Torres, J.; Berlanga, J.; Fernandez, C.; Llorente, A., Acute myeloid leukemia with MLL rearrangements: clinicobiological features, prognostic impact and value of flow cytometry in the detection of residual leukemic cells. Leukemia 2003, 17 (1), 76–82.

11. Winters, A. C.; Bernt, K. M., MLL-rearranged leukemias—an update on science and clinical approaches. Frontiers in pediatrics 2017, 5, 4.

12. Cozzio, A.; Passegué, E.; Ayton, P. M.; Karsunky, H.; Cleary, M. L.; Weissman, I. L., Similar MLL-associated leukemias arising from self-renewing stem cells and short-lived myeloid progenitors. Genes & development 2003, 17 (24), 3029–3035.

13. DiMartino, J. F.; Cleary, M. L., Mll rearrangements in haematological malignancies: lessons from clinical and biological studies. Br. J. Haematol. 1999, 106 (3), 614–626.

14. Meyer, C.; Burmeister, T.; Gröger, D.; Tsaur, G.; Fechina, L.; Renneville, A.; Sutton, R.; Venn, N.; Emerenciano, M.; Pombo-de-Oliveira, M., The MLL recombinome of acute leukemias in 2017. Leukemia 2018, 32 (2), 273–284.

15. Chan, A. K.; Chen, C.-W., Rewiring the epigenetic networks in MLL-rearranged leukemias: epigenetic dysregulation and pharmacological interventions. Frontiers in Cell and Developmental Biology 2019, 7, 81.

16. Zhao, D.; Li, Y.; Xiong, X.; Chen, Z.; Li, H., YEATS domain—A histone acylation reader in health and disease. Journal of Molecular Biology 2017, 429 (13), 1994–2002.

17. Li, Y.; Wen, H.; Xi, Y.; Tanaka, K.; Wang, H.; Peng, D.; Ren, Y.; Jin, Q.; Dent, S. Y.; Li, W., AF9 YEATS domain links histone acetylation to DOT1L-mediated H3K79 methylation. Cell 2014, 159 (3), 558–571.

18. Li, Y.; Sabari, B. R.; Panchenko, T.; Wen, H.; Zhao, D.; Guan, H.; Wan, L.; Huang, H.; Tang, Z.; Zhao, Y., Molecular coupling of histone crotonylation and active transcription by AF9 YEATS domain. Molecular cell 2016, 62 (2), 181–193.

19. Andrews, F. H.; Shanle, E. K.; Strahl, B. D.; Kutateladze, T. G., The essential role of acetyllysine binding by the YEATS domain in transcriptional regulation. Transcription 2016, 7 (1), 14–20.

20. Li, X.; Liu, S.; Li, X.; Li, X. D., YEATS Domains as Novel Epigenetic Readers: Structures, Functions, and Inhibitor Development. ACS Chemical Biology 2022.

21. Smith, E.; Lin, C.; Shilatifard, A., The super elongation complex (SEC) and MLL in development and disease. Genes & development 2011, 25 (7), 661–672.

22. Mohan, M.; Herz, H.-M.; Takahashi, Y.-H.; Lin, C.; Lai, K. C.; Zhang, Y.; Washburn, M. P.; Florens, L.; Shilatifard, A., Linking H3K79 trimethylation to Wnt signaling through a novel Dot1-containing complex (DotCom). Genes & development 2010, 24 (6), 574–589.

23. Nguyen, A. T.; Taranova, O.; He, J.; Zhang, Y., DOT1L, the H3K79 methyltransferase, is required for MLL-AF9–mediated leukemogenesis. Blood, The Journal of the American Society of Hematology 2011, 117 (25), 6912–6922.

24. Mohan, M.; Lin, C.; Guest, E.; Shilatifard, A., Licensed to elongate: a molecular mechanism for MLL-based leukaemogenesis. Nature reviews Cancer 2010, 10 (10), 721–728.

25. Erb, M. A.; Scott, T. G.; Li, B. E.; Xie, H.; Paulk, J.; Seo, H.-S.; Souza, A.; Roberts, J. M.; Dastjerdi, S.; Buckley, D. L., Transcription control by the ENL YEATS domain in acute leukaemia. Nature 2017, 543 (7644), 270–274.

26. Wan, L.; Wen, H.; Li, Y.; Lyu, J.; Xi, Y.; Hoshii, T.; Joseph, J. K.; Wang, X.; Loh, Y.-H. E.; Erb, M. A., ENL links histone acetylation to oncogenic gene expression in acute myeloid leukaemia. Nature 2017, 543 (7644), 265–269.

27. Asiaban, J. N.; Milosevich, N.; Chen, E.; Bishop, T. R.; Wang, J.; Zhang, Y.; Ackerman, C. J.; Hampton, E. N.; Young, T. S.; Hull, M. V., Cell-based ligand discovery for the ENL YEATS domain. ACS chemical biology 2020, 15 (4), 895–903.

28. Heidenreich, D.; Moustakim, M.; Schmidt, J.; Merk, D.; Brennan, P. E.; Fedorov, O.; Chaikuad, A.; Knapp, S., Structure-Based Approach toward Identification of Inhibitory Fragments for Eleven-Nineteen-Leukemia Protein (ENL). J. Med. Chem. 2018, 61 (23), 10929–10934.

29. Li, X.; Li, X. M.; Jiang, Y.; Liu, Z.; Cui, Y.; Fung, K. Y.; van der Beelen, S. H. E.; Tian, G.; Wan, L.; Shi, X.; Allis, C. D.; Li, H.; Li, Y.; Li, X. D., Structure-guided development of YEATS domain inhibitors by targeting pi-pi-pi stacking. Nat. Chem. Biol. 2018, 14 (12), 1140–1149.

30. Moustakim, M.; Christott, T.; Monteiro, O. P.; Bennett, J.; Giroud, C.; Ward, J.; Rogers, C. M.; Smith, P.; Panagakou, I.; Diaz-Saez, L.; Felce, S. L.; Gamble, V.; Gileadi, C.; Halidi, N.; Heidenreich, D.; Chaikuad, A.; Knapp, S.; Huber, K. V. M.; Farnie, G.; Heer, J.; Manevski, N.; Poda, G.; Al-Awar, R.; Dixon, D. J.; Brennan, P. E.; Fedorov, O., Discovery of an MLLT1/3 YEATS Domain Chemical Probe. Angew. Chem. Int. Ed. 2018, 57 (50), 16302–16307.

31. Ma, X. R.; Xu, L.; Xu, S.; Klein, B. J.; Wang, H.; Das, S.; Li, K.; Yang, K. S.; Sohail, S.; Chapman, A.; Kutateladze, T. G.; Shi, X.; Liu, W. R.; Wen, H., Discovery of Selective Small-Molecule Inhibitors for the ENL YEATS Domain. J. Med. Chem. 2021, 64 (15), 10997–11013.

32. Garnar-Wortzel, L.; Bishop, T. R.; Kitamura, S.; Milosevich, N.; Asiaban, J. N.; Zhang, X.; Zheng, Q.; Chen, E.; Ramos, A. R.; Ackerman, C. J., Chemical inhibition of ENL/AF9 YEATS domains in acute leukemia. ACS central science 2021, 7 (5), 815–830.

33. Wu, P.; Brand, L., Resonance energy transfer: methods and applications. Analytical biochemistry 1994, 218 (1), 1–13.

34. Dacres, H.; Michie, M.; Wang, J.; Pfleger, K. D.; Trowell, S. C., Effect of enhanced Renilla luciferase and fluorescent protein variants on the Förster distance of Bioluminescence resonance energy transfer (BRET). Biochemical and biophysical research communications 2012, 425 (3), 625–629.

35. Pfleger, K. D.; Eidne, K. A., Illuminating insights into protein-protein interactions using bioluminescence resonance energy transfer (BRET). Nature methods 2006, 3 (3), 165–174.

36. Fraga, H., Firefly luminescence: a historical perspective and recent developments. Photochemical & Photobiological Sciences 2008, 7 (2), 146–158.

37. Lorenz, W.; Cormier, M.; O’Kane, D.; Hua, D.; Escher, A.; Szalay, A., Expression of the Renilla reniformis luciferase gene in mammalian cells. Journal of bioluminescence and chemiluminescence 1996, 11 (1), 31–37.

38. Kocan, M.; See, H. B.; Seeber, R. M.; Eidne, K. A.; Pfleger, K. D., Demonstration of improvements to the bioluminescence resonance energy transfer (BRET) technology for the monitoring of G protein–coupled receptors in live cells. SLAS Discovery 2008, 13 (9), 888–898.

39. Hall, M. P.; Unch, J.; Binkowski, B. F.; Valley, M. P.; Butler, B. L.; Wood, M. G.; Otto, P.; Zimmerman, K.; Vidugiris, G.; Machleidt, T., Engineered luciferase reporter from a deep sea shrimp utilizing a novel imidazopyrazinone substrate. ACS chemical biology 2012, 7 (11), 1848–1857.

40. Robers, M. B.; Dart, M. L.; Woodroofe, C. C.; Zimprich, C. A.; Kirkland, T. A.; Machleidt, T.; Kupcho, K. R.; Levin, S.; Hartnett, J. R.; Zimmerman, K., Target engagement and drug residence time can be observed in living cells with BRET. Nature communications 2015, 6 (1), 1–10.

41. Gnatzy, M. T.; Geiger, T. M.; Kuehn, A.; Gutfreund, N.; Walz, M.; Kolos, J. M.; Hausch, F., Development of NanoBRET-Binding Assays for FKBP-Ligand Profiling in Living Cells. ChemBioChem 2021, 22 (13), 2257–2261.

42. Garnar-Wortzel, L.; Bishop, T. R.; Kitamura, S.; Milosevich, N.; Asiaban, J. N.; Zhang, X.; Zheng, Q.; Chen, E.; Ramos, A. R.; Ackerman, C. J.; Hampton, E. N.; Chatterjee, A. K.; Young, T. S.; Hull, M. V.; Sharpless, K. B.; Cravatt, B. F.; Wolan, D. W.; Erb, M. A., Chemical Inhibition of ENL/AF9 YEATS Domains in Acute Leukemia. ACS Cent. Sci. 2021, 7 (5), 815–830.

43. Lazareno, S.; Birdsall, N., Estimation of competitive antagonist affinity from functional inhibition curves using the Gaddum, Schild and Cheng-Prusoff equations. British journal of pharmacology 1993, 109 (4), 1110.

44. Li, A. P., Screening for human ADME/Tox drug properties in drug discovery. Drug discovery today 2001, 6 (7), 357–366.

45. Eddershaw, P. J.; Beresford, A. P.; Bayliss, M. K., ADME/PK as part of a rational approach to drug discovery. Drug discovery today 2000, 5 (9), 409–414.

46. Masimirembwa, C. M.; Bredberg, U.; Andersson, T. B., Metabolic stability for drug discovery and development. Clinical pharmacokinetics 2003, 42 (6), 515–528.

47. Mandagere, A. K.; Thompson, T. N.; Hwang, K.-K., Graphical model for estimating oral bioavailability of drugs in humans and other species from their Caco-2 permeability and in vitro liver enzyme metabolic stability rates. Journal of medicinal chemistry 2002, 45 (2), 304–311.

48. Houston, J. B., Utility of in vitro drug metabolism data in predicting in vivo metabolic clearance. Biochemical pharmacology 1994, 47 (9), 1469–1479.

49. Ito, K.; Ogihara, K.; Kanamitsu, S.-i.; Itoh, T., Prediction of the in vivo interaction between midazolam and macrolides based on in vitro studies using human liver microsomes. Drug Metabolism and Disposition 2003, 31 (7), 945–954.

