## Supplementary Material for "The Prioritization of Eleven-Nineteen-Leukemia Inhibitors as Potential Drug Candidates to Treat Acute Myeloid Leukemia"

for

**Materials.** All reagents and solvents for synthesis were purchased from commercial sources and used without purification. All glassware were flame-dried prior to use. Thin-layer chromatography (TLC) was carried out on aluminum plates coated with 60 F254 silica gel. TLC plates were visualized under UV light (254 or 365 nm) or stained with 5% phosphomolybdic acid. Intracellular TE Nano-Glo® Substrate/Inhibitor (N2161) for NanoBRET assay was purchased from Promega. Plasmid pCDH-EF1-Nluc (73024) was from Addgene. Nluc Luciferase Antibody (MAB10026-SP) was from Biocompare. Frozen human plasma was purchased from Gulf Coast Regional Blood Center. Pooled human liver microsome (1910096) was from Xenotech. Human cell lines HEK 293T/17, MV4-11, MOLM-13, JURKAT cells were purchased from ATCC. RPMI 1640 medium (11875093), DMEM (11965092), Opti-MEM (2491765), RPMI 1640 without phenol red (2420200), Heat inactivated Fetal Bovine Serum (FBS) (16140071) and puromycin (2149042) were purchased from Gibco. IMDM (30-2005) was purchased from ATCC. Non-heat inactivated FBS (SH30071.03) was from Cytiva. Penicillin/streptomycin (17-602E) was from Lonza. Cell Cycle Analysis Kit (ab287852) and Cell Counting Kit-8 (ab228554) were from Abcam. EndoFree Plasmid Midi Kit (D6915-03) was from Omega Bio-tek Inc. Lenti-X GoStix Plus Kit (631280) was provided by Takara Inc.

**Instruments.** Normal phase column chromatography was carried out using a Yamazen Smart Flash AKROS system. NMR spectra were recorded on a Bruker AVANCE Neo 400 MHz or Varian INOVA 300 MHz spectrometer in specified deuterated solvents. High-resolution electrospray ionization mass spectrometry (HRMS-ESI) was carried out on a Thermo Scientific Q Exactive Focus system. NanobBRET assay and cell viability were carried on BioTek SYNERGY neo2 multi-mode reader. Cell cycle was analyzed by Beckman Coulter Cytoflex.

**Cell Lines and Culture Condition.** MV4-11 (IMDM supplemented with 10% non-heat inactivated FBS, penicillin, and streptomycin), MOLM-13 (RPMI 1640 supplemented with 10% heat inactivated FBS, penicillin, and streptomycin), JURKAT (RPMI 1640 supplemented with 10% heat inactivated FBS, penicillin, and streptomycin), HEK 293T (DMEM supplemented with 10% heat inactivated FBS, penicillin, and streptomycin) were maintained in a humidified 37 °C incubator with 5% $CO_2$. Cells at logarithmic growth phase were used for following experiments.

**Generation of stable Nluc-ENL YEATS expressed HEK293T cell line.** cDNA encoding ENL YEATS domain (aa 1-148) was into a Nluc containing plasmid pCDH-EF1-Nluc after removing the Nluc TAA STOP to afford pCDH-EF1-Nluc-ENL YEATS and then Nluc-ENL YEATS was cloned into pLVx-IRES-Puro to give PLVx- Nluc-ENL YEATS. The plasmid for lentivirus packaging was prepared using the EndoFree Plasmid Midi Kits from according to the manufacturer's protocol. For packaging Nluc-ENL YEATS lentivirus particles, HEK293T/17 cells were cultured in 10 $cm^2$ dishes to 70-80% confluency and then co-transfected with three plasmids pLVx- Nluc-ENL YEATS, psPAX2 and PMD2.G using polyethyleneimine as described previously.[1] The transfected cells were cultured for 2 days and the supernatants were collected to isolate viral particles. Additional medium (10 mL) was provided to transfected cells for growing one more day for subsequent supernatants collection. The collected supernatants were centrifuged at 1500 rpm at 4 °C for 5 min to remove cell debris, residual supernatants were then filtered through a 0.45 μm membrane, and then centrifuged at 30,000× g at 4 °C for 4 h to precipitate viral particles. After that, the supernatants were discarded carefully, and the lentiviral pellets were resuspended in serum-free DMEM medium (200 μL) and stored at -80 °C. Successful production of lentivirus and tittered collected lentiviral particle were confirmed and detected using Lenti-X GoStix Plus Kits provided by Takara Inc. following standard protocols. To transduce HEK293T cells, we incubated them with suspended lentiviral solutions (MOI 5-10) for 24 h with 4 μg/mL polybrene in serum-free DMEM medium before replacing fresh DMEM. After 2 days, cells were selected for stable expression of Nluc-ENL YEATS using puromycin (1μg/mL) for 2 weeks. The expression of Nluc-ENL YEATS was then confirmed by immunoblotting using Nluc luciferase antibody.

**Chemical synthesis**

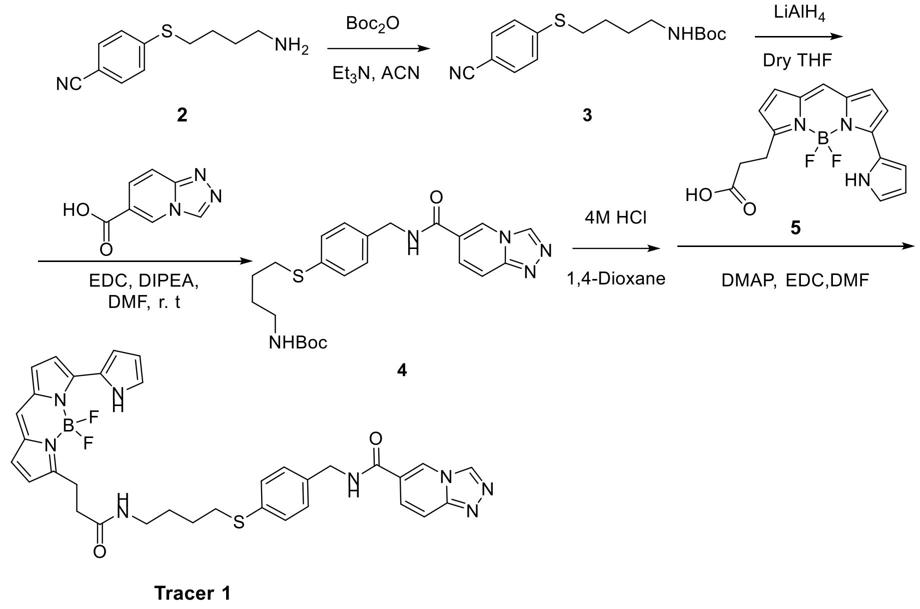


**Scheme S1**. The synthesis of **Tracer 1**.

Compound **3.** Compound **2** (2.43 mmol, 500 mg) was dissolved in ACN (25 mL) under nitrogen atmosphere. A solution of Boc2O (2.67 mmol, 581 mg) in dry ACN (25 mL) was added dropwise at 0°C under stirring. The solution was stirred overnight at room temperature. The solvent was removed under reduced pressure and the residue was then purified by column chromatography (silica gel, 10% EtOAc/n-Hexane as the eluent) to yield **3** as a white solid (578 mg, 78%). $^1$H NMR (400 MHz, Chloroform-d) δ 7.52 (d, J = 8.6 Hz, 1H), 7.29 (d, J = 8.6 Hz, 1H), 4.53 (s, 1H), 3.15 (q, J = 5.9 Hz, 2H), 2.99 (t, J = 7.1 Hz, 1H), 1.76-1.70 (m, 2H), 1.68-1.61 (m, 2H), 1.43 (s, 5H).

Compound **4.** $LiAlH_4$ (2.08 mmol, 77 mg) was suspended in dry THF (10 mL) under nitrogen atmosphere and was cooled under 0 °C. The solution of compound **3** (1.90 mmol, 578 mg) in dry THF (10 mL) was added dropwise at 0°C under stirring. The solution was stirred 4 h at room temperature. Water (3 mL) was added dropwise, followed by 2 M aqueous NaOH solution (3 mL) and then, water (3 mL). The precipitate was filtered and washed with THF. The combined filtrate was then dried with anhydrous $Na_2SO_4$ and evaporated in vacuo. The residue was directly dissolved with DMF (10 mL), then 1,2,4-triazolo [4,3-a] pyridine-6-carboxylic acid (2 mmol, 326 mg), DIPEA (4 mmol, 516 mg) and EDCI (2.4 mmol, 460 mg) were added. The resulting solution was stirred at room temperature overnight. Then, the solution was diluted with EtOAc (40 mL) and washed with saturated $NaHCO_3$ solution (2 × 50 mL), 1 M HCl (2 × 50 mL), and saturated brine (50 mL). The organic layers were then dried with anhydrous $Na_2SO_4$ and then concentrated. The residue was purified by column chromatography (silica gel, 10% MeOH/DCM as the eluent) to yield **4** as a light-yellow solid (446 mg, 54%). $^1H$ NMR (400 MHz, Chloroform-d) δ 8.91 (d, J = 1.6 Hz, 1H), 8.77 (s, 1H), 7.59 (t, J = 8.2 Hz, 2H), 7.47-7.39 (m, 1H), 7.17 (s, 4H), 4.54 (d, J = 5.5 Hz, 3H), 4.24 - 4.15 (m, 5H), 3.07-2.93 (m, 2H), 2.82 (t, J = 6.7 Hz, 2H). HRMS (ESI) calculated for $C_{23}H_{29}N_5O_3S$ $[M+H]^+$ 456.2064, found 456.2055.

**Tracer 1**. To a solution of **4** (0.1 mmol, 46 mg) in 5 mL of 1,4-dioxane was added 5 mL of 4 M HCl solution in 1,4-dioxane. The resulting solution was stirred at room temperature for 2 h. Then, the reaction mixture was concentrated to dryness in vacuo. The residue was directly dissolved in anhydrous DMF (2 mL), compound **5** (0.10 mmol, 36 mg) and DMAP (0.20 mmol, 24 mg) were added. The mixture was cooled to 0°C and EDC (0.15 mmol, 29 mg) was added. The resulting mixture was then stirred at room temperature for 36 h. The reaction mixture was purified by column chromatography (silica gel, 5% methanol/DCM as the eluent) to yield fluorescence tracer **1** as a purple solid (6.2 mg, 9.3 %). 1H NMR (400 MHz, Chloroform-d) δ 10.30 (s, 1H), 8.74 (d, J = 0.8 Hz, 1H), 8.71 (t, J = 1.4 Hz, 1H), 7.68-7.63 (m, 1H), 7.54-7.50 (m, 1H), 7.40-7.34 (m, 2H), 7.29 (s, 1H), 7.23-7.21 (m, 1H), 7.12 (d, J = 8.2 Hz, 1H), 7.08-7.02 (m, 2H), 6.94 (s, 1H), 6.89 (d, J = 4.7 Hz, 1H), 6.81 (d, J = 8.2 Hz, 1H), 6.75 (d, J = 4.0 Hz, 1H), 6.43-6.36 (m, 2H), 5.90 (t, J = 6.1 Hz, 1H), 4.56 (dd, J = 5.4, 3.5 Hz, 2H), 4.43 (d, J = 4.1 Hz, 2H), 3.31-3.18 (m, 2H), 2.97-2.86 (m, 2H), 2.74 -2.55 (m, 2H), 1.61 (d, J = 6.8 Hz, 4H). HRMS (ESI) calculated for $C_{34}H_{32}BF_2N_8O_2S$ $[M-H]^-$ 665.2425, found 665.2452.

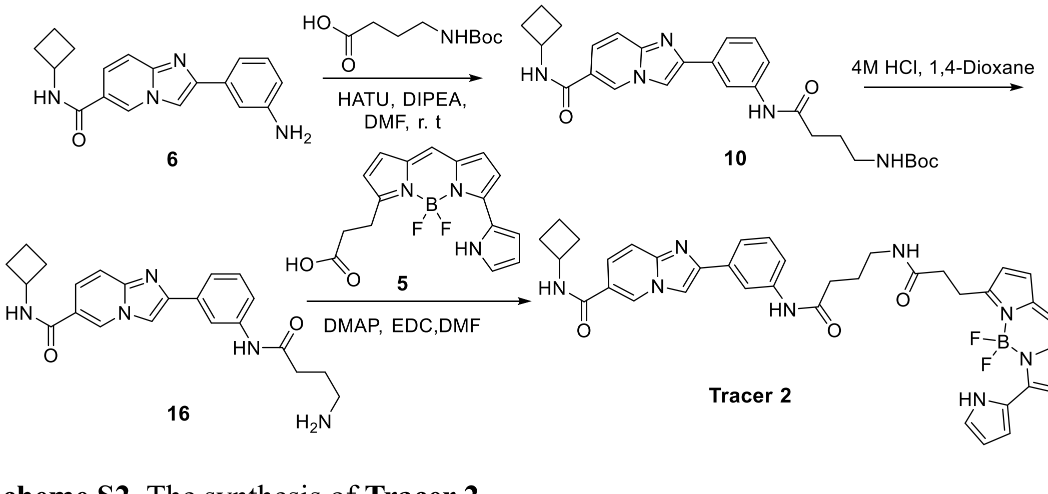


**Scheme S2**. The synthesis of **Tracer 2.**

Compound **6**[2] and **5**[3] was synthesized according to reported literature.

Compound **10.** To a solution of N-(tert-butoxycarbonyl)-4-aminobutyric acid (1.57 mmol, 225 mg) in anhydrous DMF (15 mL) was added DIPEA (3.15 mmol, 407 mg) for 30 min stir at r.t, then HATU (1.25 mmol, 475 mg) and compound **6** (2.1 mmol, 530 mg) were added. The resulting solution was then stirred at room temperature overnight. The reaction mixture was then diluted with EtOAc (50 mL) and then washed with saturated $NaHCO_3$ solution (2 × 50 mL), 0.5 M HCl (2 × 10 mL), and brine (50 mL). The organic layers were dried with anhydrous $Na_2SO_4$ and concentrated in vacuo. The residue was then purified by column chromatography (silica gel, 5% methanol/DCM as the eluent) to yield **10** as a white solid (420 mg, 68%). $^1$H NMR (400 MHz, DMSO-$d_6$): δ 9.99 (s, 1H), 9.07 (dd, *J* = 1.8, 1.0 Hz, 1H), 8.73 (d, *J* = 7.5 Hz, 1H), 8.43 (s, 1H), 8.27 (t, *J* = 1.9 Hz, 1H), 7.68 (dd, *J* = 9.5, 1.8 Hz, 1H), 7.64-7.56 (m, 3H), 7.36 (t, *J* = 7.9 Hz, 1H), 6.85 (t, *J* = 5.8 Hz, 1H), 2.98 (q, *J* = 6.6 Hz, 2H), 2.32 (t, *J* = 7.5 Hz, 2H), 2.24 (dd, *J* = 8.7, 2.9 Hz, 2H), 2.15-2.02 (m, 2H), 1.75-1.65 (m, 4H), 1.38 (s, 9H). 13C NMR (101 MHz, DMSO-$d_6$) δ 171.47, 163.60, 156.09, 146.03, 145.37, 140.25, 134.39, 129.53, 128.69, 124.02, 120.91, 120.37, 119.15, 116.88, 116.11, 110.50, 77.93, 45.09, 30.56, 28.74, 26.05, 15.23. HRMS (ESI) calculated for $C_{27}H_{34}N_5O_4$ $[M+H]^+$ 492.2606, found 462.2596.

Compound **16.** To a solution of **10** (0.5 mmol, 245 mg) in 5 mL of 1,4-dioxane was added 10 mL of 4 M HCl solution in 1,4-dioxane. The resulting solution was stirred at room temperature for 2 h. Then, the reaction mixture was concentrated to dryness in vacuo to yield **16** as a brown solid (178 mg, 91%).$^1$H NMR (400 MHz, DMSO-$d_6$) δ 10.42 (s, 1H), 9.34 (s, 1H), 9.09 (d, *J* = 7.4 Hz, 1H), 8.68 (s, 1H), 8.33 (t, *J* = 1.9 Hz, 1H), 8.16 (d, *J* = 9.3 Hz, 1H), 8.01 (s, 3H), 7.90 (d, *J* = 9.4 Hz, 1H), 7.71 (dt, *J* = 7.8, 1.3 Hz, 1H), 7.65 (dt, *J* = 8.2, 1.3 Hz, 1H), 7.50 (t, *J* = 7.9 Hz, 1H), 4.47 (p, *J* = 8.1 Hz, 1H), 2.87 (h, *J* = 5.9 Hz, 2H), 2.32-2.22 (m, 2H), 2.19-2.05 (m, 2H), 1.91 (p, *J* = 7.3 Hz, 2H), 1.72 (tt, *J* = 10.5, 5.1 Hz, 2H). $^{13}$C NMR (101 MHz, DMSO-$d_6$) δ 171.27, 162.14, 141.72, 140.56, 137.85, 131.16, 130.29, 124.25, 117.18, 112.40, 112.07, 72.62, 70.98, 66.82, 63.26, 60.64, 45.32, 38.80, 33.50, 30.36, 23.47, 15.30. HRMS (ESI) calculated for $C_{22}H_{26}N_5O_4$ $[M+H]^+$ 392.2082, found 392.2075.

**Tracer 2**. To a solution of **16** (0.10 mmol, 40 mg) in anhydrous DMF (2 mL) was added compound **5** (0.10 mmol, 36 mg) and DMAP (0.20 mmol, 24 mg). The mixture was cooled to 0°C and EDC (0.15 mmol, 29 mg) was added. The resulting mixture was then stirred at room temperature for 36 h. The reaction mixture was purified by column chromatography (silica gel, 5% methanol/DCM as the eluent) to yield fluorescence tracer **2** as a purple solid (6.8 mg, 8.9 %). $^1$H NMR (400 MHz, Chloroform-*d*) δ 10.41 (s, 1H), 8.85 - 8.72 (m, 1H), 8.68 (s, 1H), 8.15 (s, 1H), 7.91 (s, 1H), 7.72 (d, *J* = 5.4 Hz, 1H), 7.60 (t, *J* = 8.3 Hz, 3H), 7.38 (d, *J* = 9.3 Hz, 2H), 7.12 (d, *J* = 3.6 Hz, 1H), 7.01 (d, *J* = 4.6 Hz, 1H), 6.96 (d, *J* = 4.4 Hz, 1H), 6.82 (dd, *J* = 8.3, 4.2 Hz, 2), 6.35-6.33 (m, 1H), 6.30-6.24 (m, 2H), 6.08 (s, 1H), 4.64-4.55 (m, 1H), 3.39-3.35 (m, 4H), 2.74 (t, *J* = 7.4 Hz, 2H), 2.49-2.41 (m, 2H), 2.32-2.24 (m, 2H), 2.00-1.94 (m, 2H), 1.87-1.77 (m, 4H). HRMS (ESI) calculated for $C_{38}H_{38}BF_2N_8O_3$ $[M+H]^+$ 703.3123, found 703.3109.

**NanoBRET assay**

All NanoBRET measurements were based on the protocols described in previous publications and modified as described below.[4]

**Apparent Tracer Affinity.** Approximately 18000 Nluc-ENL YEATS expressed HEK293T cells (100 μL) were seeded into a white non-binding 96-well assay plate and incubated overnight at 37 °C in a humidified 5% $CO_2$ atmosphere. The fluorescent tracer was dissolved in DMSO at thousand-fold the highest concentration required for the final sample. The tracer stock solution was used for a serial dilution in DMSO. Each dilution was further diluted in Opti-MEM medium to obtain the concentration required for the final sample. After removal of the medium, the cells were added with tracer in the medium (100 μL) and Opti-MEM with 0.1 % DMSO (10 μL). As a no tracer control cells were treated with 110 μL Opti-MEM with 0.1 % DMSO. As tracer + excess unlabeled group, cells were treated with tracer in the medium (100 μL) and excess unlabeled compound **6** or inhibitor **1** (880 μM, 10 μL). After incubation at 37 °C for two hours the assay plate was equilibrated at room temperature for 15 min. For BRET detection, the NanoGlo Substrate and the extracellular NanoLuc® inhibitor were diluted with Opti-MEM as described in protocol to afford detection solution. 50 μL of the detection solution was added per well and the plate was incubated for three minutes at room temperature. The donor emission was measured at 450 nm and the acceptor emission at 610 nm using BioTek SYNERGY neo2 multi-mode reader. The BRET ratio was calculated according to the following formula: BRET Ratio = $[(Acceptor_{sample}/Donor_{sample}) - (Acceptor_{no\text{-}tracer\ control}/Donor_{no\text{-}tracer\ control})] \times 1000$.

**ENL Inhibitor Competitive Assay.** Sample procedures were followed for cell seeding as described in tracer affinity assay. After removal of the medium, the cells were added with tracer in the medium (1.1 μM, 100 μL) and different concentration of test compound in 0.1 % DMSO (10 μL). Thoroughly mix plate on an orbital shaker for 15 seconds at 900 rpm. Then the cells were incubated at 37°C, 5% $CO_2$ for 2 hours for BRET ratio detection as illustrated previously.

***In Vitro* Stability in Human Plasma.** The *in* vitro stability of the test compounds was initiated by the addition of the test compounds to 90 μL of pre-warmed (37°C) human plasma yield a final concentration of 5 μM. The assays were performed in a plate shaker at 37°C and conducted in triplicate. At 0, 5, 15, 30, 60, 120 min, 400 μL acetonitrile (with internal standard Diclofenac10 μg/mL) in order to deproteinize the plasma and terminate the reaction. The samples were subjected to vortex mixing for 1 min and then centrifugation at 4°C for 20 min at 10000 rpm. The 200 μL of clear supernatants were analyzed by HPLC-MS/MS. The values represent the mean of three independent experiments. The precent of test compound remaining was determined by following formula: % remaining = (Area at $t_x$/Average area at $t_0$) × 100.

***In Vitro* Metabolic Stability in Human Liver Microsomes (HLM).** This metabolic stability measurements were based on previous publications and modified as described below.[5] Metabolic stability profile of ENL inhibitor, including in $CL_{int,\ pred}$ and *in vitro* $t_{1/2}$ was determined by the estimation of the remaining compound concentration after incubation with HLM, NADPH (cofactor), and $MgCl_2$ in a 0.1 M phosphate buffer (pH 7.4). Briefly, preincubation of 5 μM of compound was carried out in 40 μL HLM (0.5 mg/mL) in 0.1 M phosphate buffer (pH 7.4) at 37°C for 10 min to set optimal a condition for metabolic reactions. After pre incubation, NADPH (5 mM, 10 μL) or 0.1 M PB (10 μL) was then added to initiate metabolic reaction at 37°C with gentle shaking. To confirm results, the same metabolic experiment was repeated three times. At 0, 5, 15, 30, 45, 60 min, 200 μL acetonitrile (with internal standard Diclofenac10 μg/mL) in order to terminate the reaction. The samples were subjected to vortex mixing for 1 min and then centrifugation at 4°C for 10 min at 10000 rpm. Then 200 μL of clear supernatants were analyzed by HPLC-MS/MS. The precent of test compound remaining was determined by following formula: % remaining = (Area at $t_x$ /Average area at $t_0$) × 100. The half-life ($t_{1/2}$) was calculated using the slope (k) of the log-linear regression from the % remaining parent compound versus time (min) relationship: $t_{½}$ (min) = –ln 2/k. $CL_{int, pred}$ (mL/min/kg) was calculated through following formula $CLi_{nt, pred}$ = (0.693/$t_{1/2}$) × (1/ (microsomal protein concentration (0.5 mg/mL)) × Scaling Factor (1254.16 for human liver microsome).

**Cell Viability Assay.** Approximately 10000 JURKAT, MOLM-13, MV4-11 and HEK293T cells in RPMI 1640(no phenol red) (100 μL) were added to a 96-well plate and treated with DMSO or compounds at indicated concentrations for 72 h. Cell viability was measured using the CCK-8 kit according to the manufacturer's instructions. The plates were placed on a microplate reader at ambient temperature and the absorbance at 460 nm of each well was taken. The average absorbance of the blank wells, which did not contain the cells, was subtracted from the reading of the other wells. The cell viability was then determined by the equation: % Viability = [Σ ($A_i$/$A_{control}$ × 100)]/n, where Ai is the absorbance of the $i_{th}$ data (i = 1, 2, …, n), $A_{control}$ is the absorbance of the control wells, in which DMSO was added, and n is the number of data points.

**Anti-Proliferation Assay.** Cell proliferation assays were carried out in 96-well tissue culture plates at 20000 cells/well for all cell lines except in 200 μL with compounds added as 1:1000 dilutions of DMSO stocks in triplicate. Culture density was determined every 3-4 days using the Countess automated cell counter, after which 20000 live cells were reseeded in fresh media and compound. The cumulative cell count was achieved by back calculation.

**Cell Cycle Analysis.** For cell cycle analysis, 500k cells were cultured at 2 mL/well in a 6-well plate treated in a ratio of 1:1000 with compound stocks in DMSO or 1:1000 with DMSO in triplicate. Cells cycle staining was performed by using cell cycle analysis kit and treated according to the protocol provided. The signal was analyzed by flow cytometry and the cytometry results were analyzed by CytExpert software.

**Cellular Thermal Shift Assay.** MOLM-13 and MV4-11 were incubated with 10 μM of compound **13** for 3 h and 6 h, respectively. The cells were then collected and washed with PBS 3 times. The cell pellets were resuspended in PBS-containing protease inhibitors and aliquoted into PCR microtubes (approximately 3 million cells in 54 μL). Cells were heated at indicated temperatures for 3 min in a thermal cycler (Bio-Rad) and then incubated at room temperature for 2 min. A total of 6 μL of 10× cell lysis buffer

(8% NP-40, 50% glycerol, and 10 mM dithiothreitol) was added to each sample before subjecting to three freeze-thaw cycles by liquid nitrogen and 37 °C water bath incubations to lyse the cells. The cell lysates were centrifuged at 13,000 rpm at 4 °C for 10 min, and the supernatants were analyzed using SDS-PAGE and western blotting.

**RNA Extraction and qRT-PCR.** Total RNA was extracted using the RNeasy plus kit (Qiagen) and reverse-transcribed using an iScript cDNA synthesis kit (Bio-Rad). Quantitative real-time PCR (qRTPCR) analyses were performed as described previously using PowerUp SYBR Green PCR Master Mix and the Bio-Rad CFX96 real-time PCR detection system. Gene expressions were calculated following normalization to β2-Microglobulin (B2M) levels using the comparative Ct (cycle threshold) method. The primer pairs are as follows. HOXA9: forward 5′-GTATAG-GGGCACCGCTTTTT-3′, reverse 5′-AATGCTGAGAATGAGAGCGG-3′. MEIS1: forward 5′-CACGCTTTTTGTGACGCTT-3′, reverse 5′ GGACAACAGCAGTG-AGCAAG-3′. MYB: forward 5’- GATGTGTGACCATGACTATG-3’, reverse 5’-GCACTGCACATCTGTTCGAT-3’, MYC: forward 5′-CACCGAGTCGTAGTCG-AGGT-3′, reverse 5′-TTT-CGGGTAGTGGAAAACCA-3′. B2M: forward 5′-AATGTCGGATGGATGAAACC-3′ reverse 5′-TAGCTGTGCTCGCGCTACT-3′.

**Pharmacokinetics Study.** Male CD-1 mice were used in the PK study. Inhibitor **13** was dissolved in a mixed solution containing 75% PEG300 and 25% D5W (5% dextrose in distilled water) and another solution containing 0.5% methyl cellulose, 0.5% Tween 80 in water at doses of 20 mg/kg for **i.v.** and **p.o.** administrations, respectively. Three mice were used for each administration, blood samples were taken via a vein at different time point up to 24 hours after dosing, collected in tubes coated with an anticoagulant, and centrifuged at 15,000g for 5 min to obtain plasma samples. Acetonitrile-containing internal standard (Labetalol, 100 ng/mL) was added to the plasma to precipitate proteins. The samples were subjected to vortex mixing for 10 min and then centrifugation at 4°C for 15 min at 3220 g. Then clear supernatants were analyzed by HPLC-MS/MS.

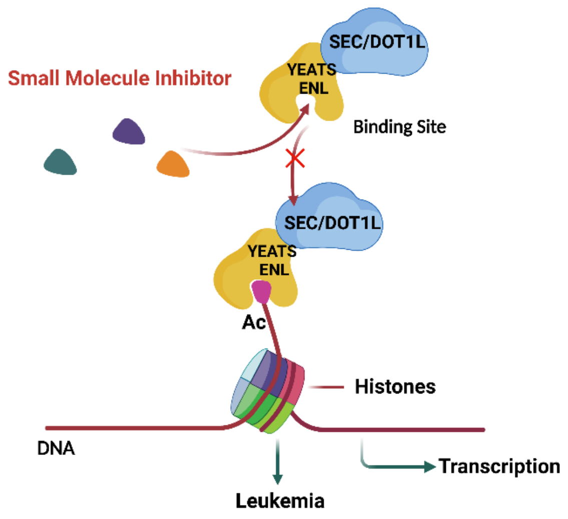


Figure S1. Disrupting the interactions between ENL YEATS domain and acetylated histones inhibits leukemia progress.

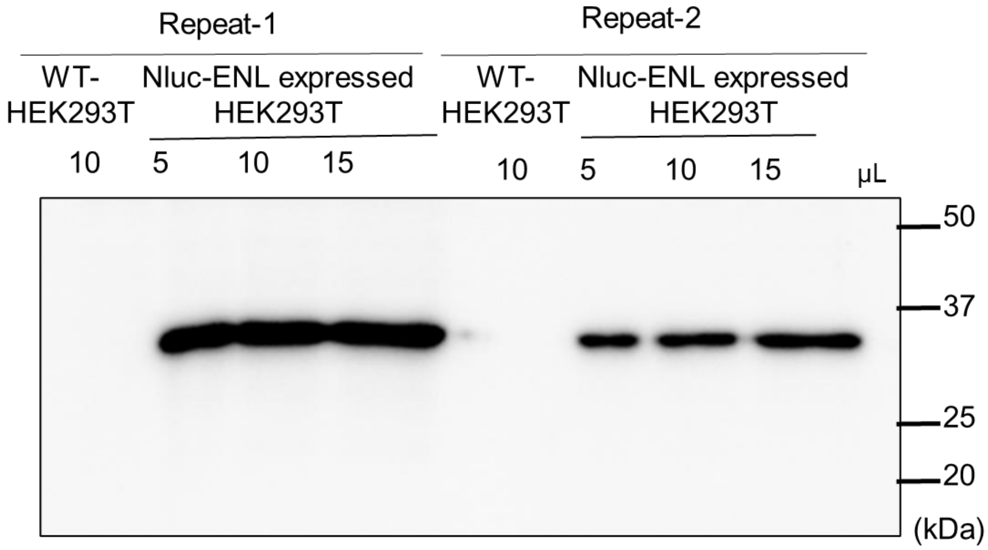


Figure S2. Expression of Nluc-ENL in Nluc-ENL expressed HEK293T and WT HEK293T cell.

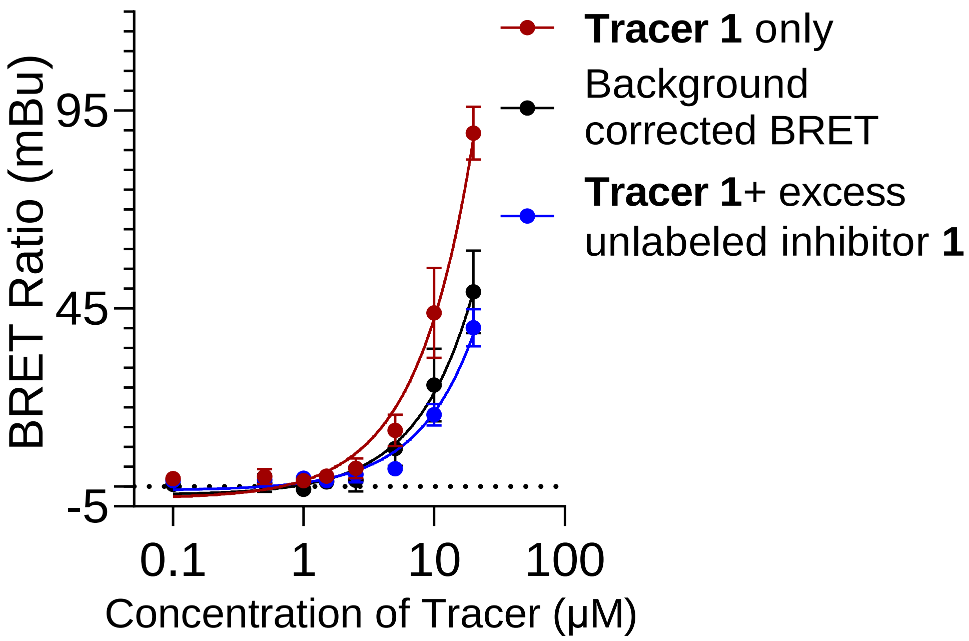


Figure S3. Apparent affinity of **Tracer 1** for NLuc-ENL YEATS fusion protein in HEK293T cells.

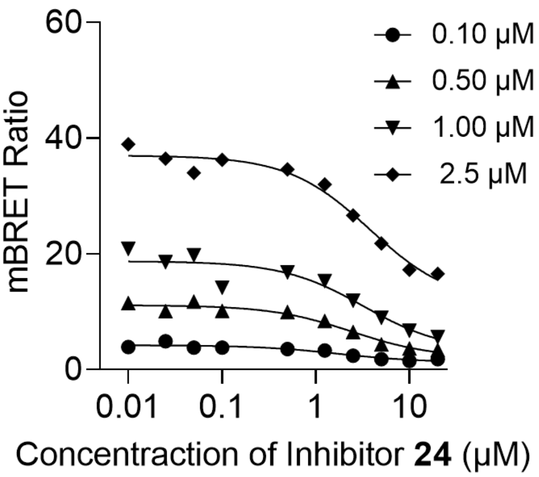


| Tracer (μM) | $IC_{50}$ (μM) |
|---|---|
| 0.10 | 1.74 |
| 0.50 | 2.43 |
| 1.00 | 3.25 |
| 2.50 | 3.78 |

Figure S4. NanoBRET curves of ENL inhibitor **24** affinities for NLuc-ENL YEATS in HEK293T cells with different concentration of **Tracer 2** and corresponding $IC_{50}$ values.

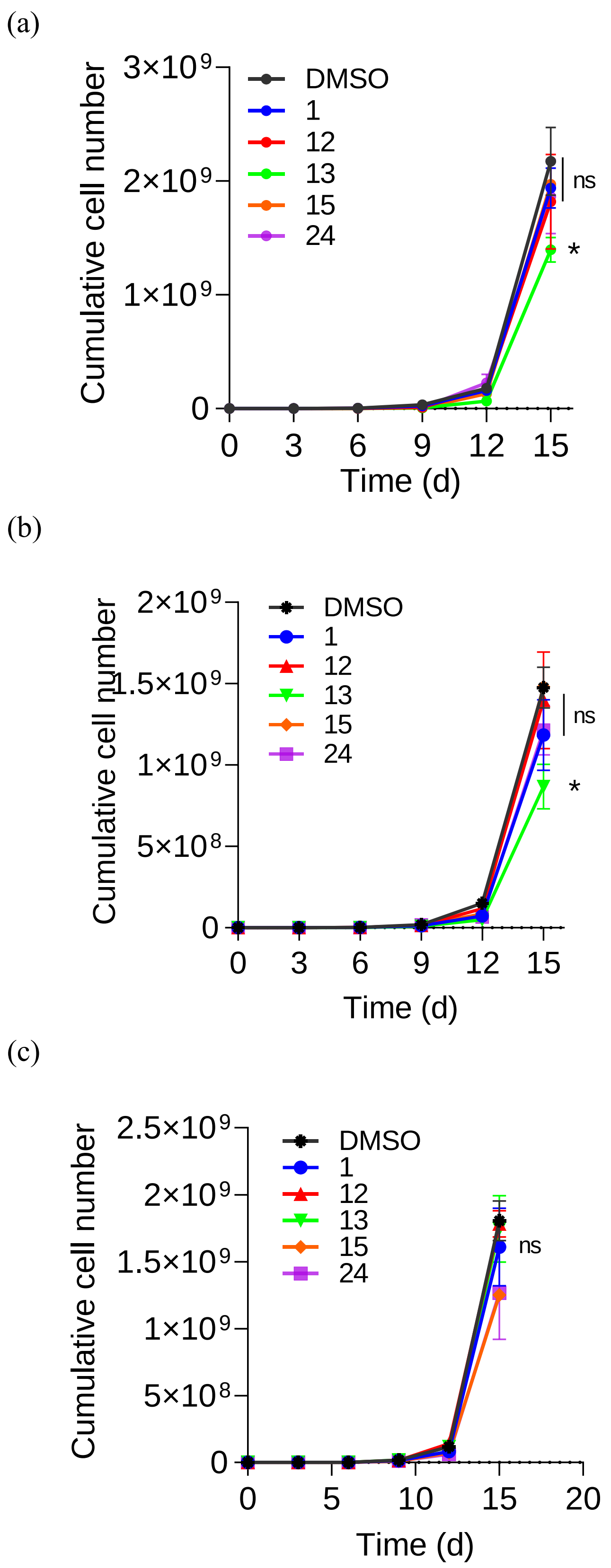


Figure S5. Proliferation of (a) MOLM-13, (b) MV4-11 and (c) Jurkat cell lines in response to ENL inhibitor in 1 μM.

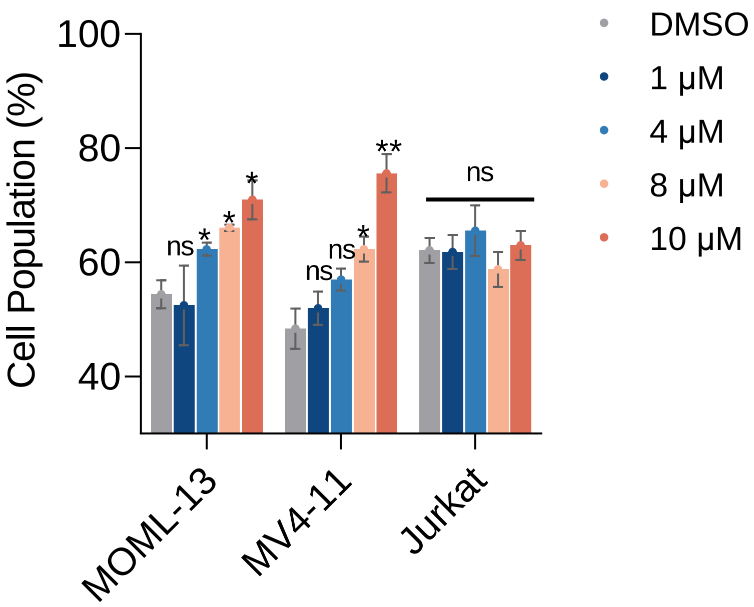


Figure S6. Comparison of percentage of cells in G1 phase in different cells after 72 h treated with different concentration of inhibitor **13**. *P < 0.05, **P < 0.01, Not significant (n.s.) P > 0.05.

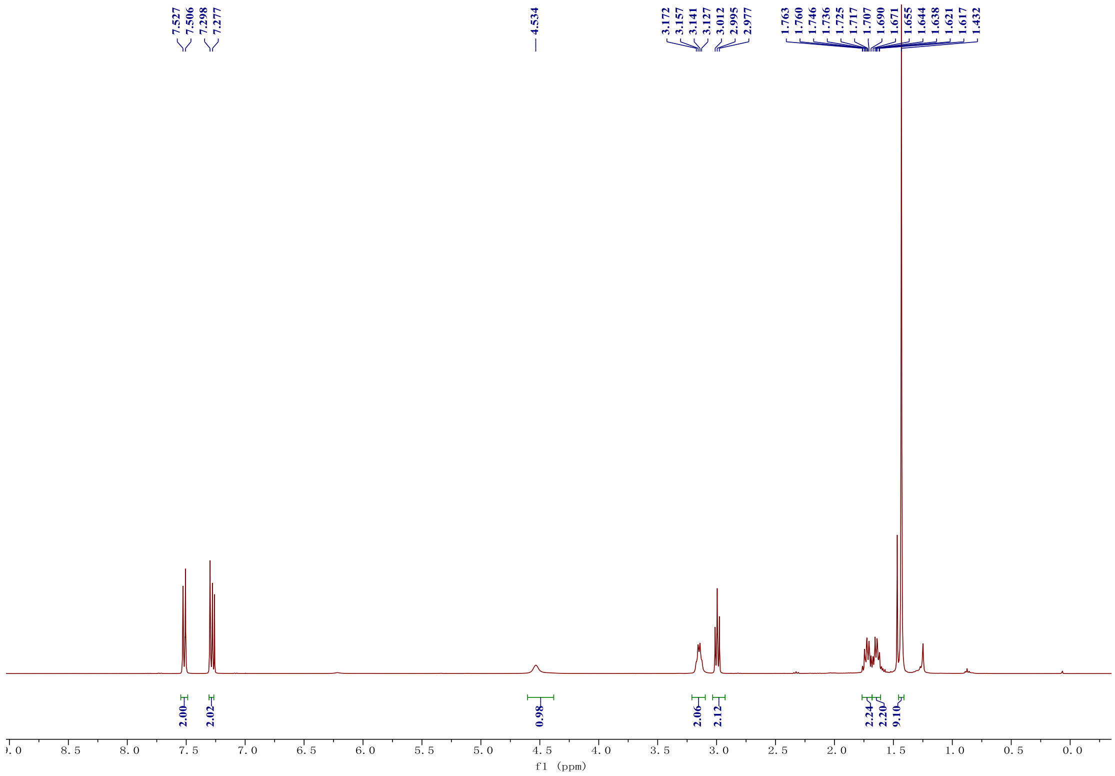


Figure S7. [1]H NMR of **3** in Chloroform-d.

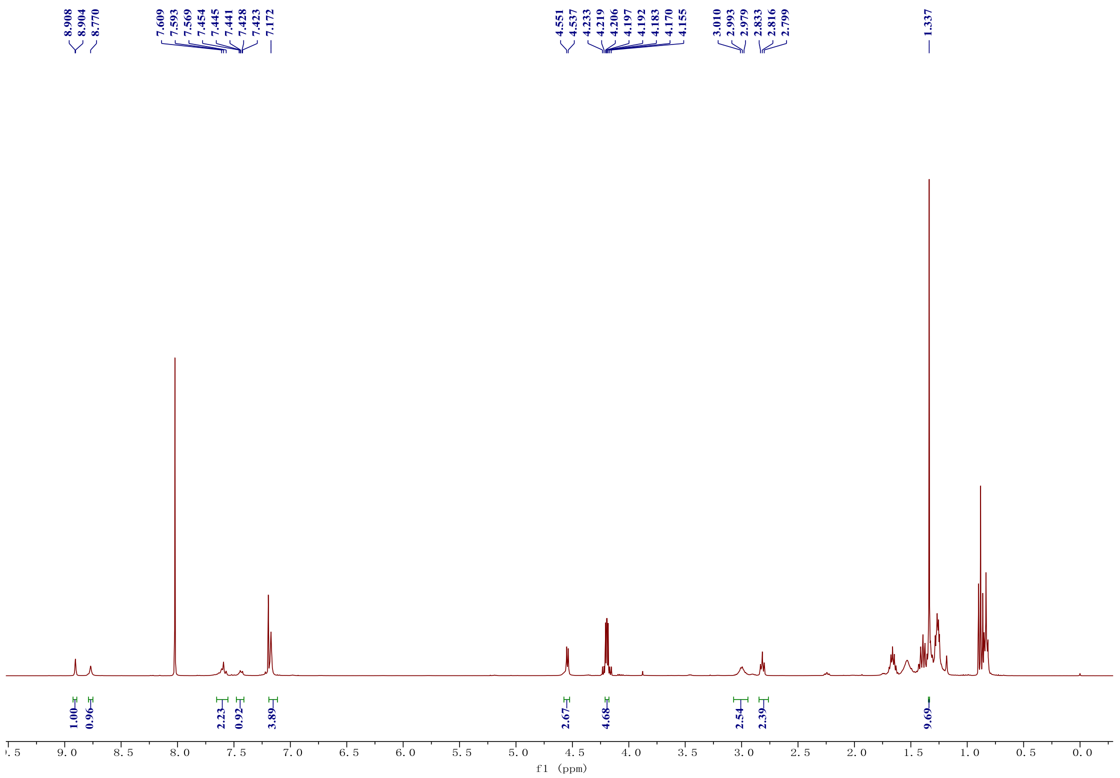


Figure S8. [1]H NMR of **4** in Chloroform-d.

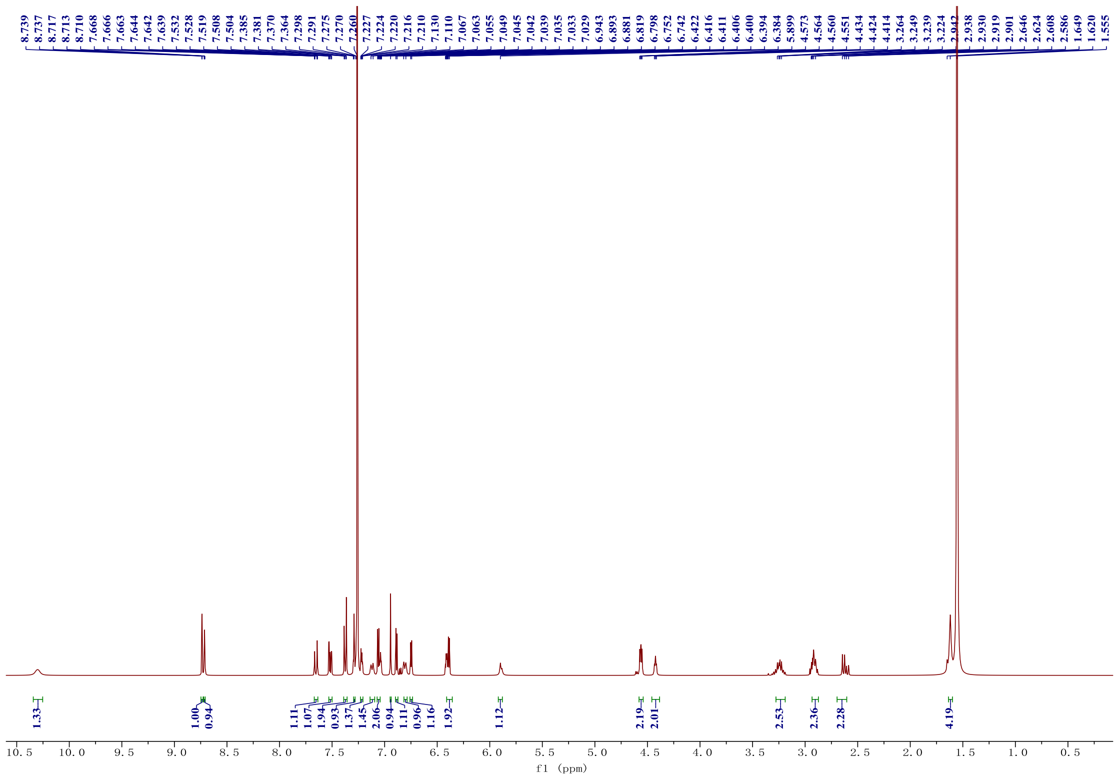

Figure S9. $^1$H NMR of **Tracer 1** in Chloroform-d.

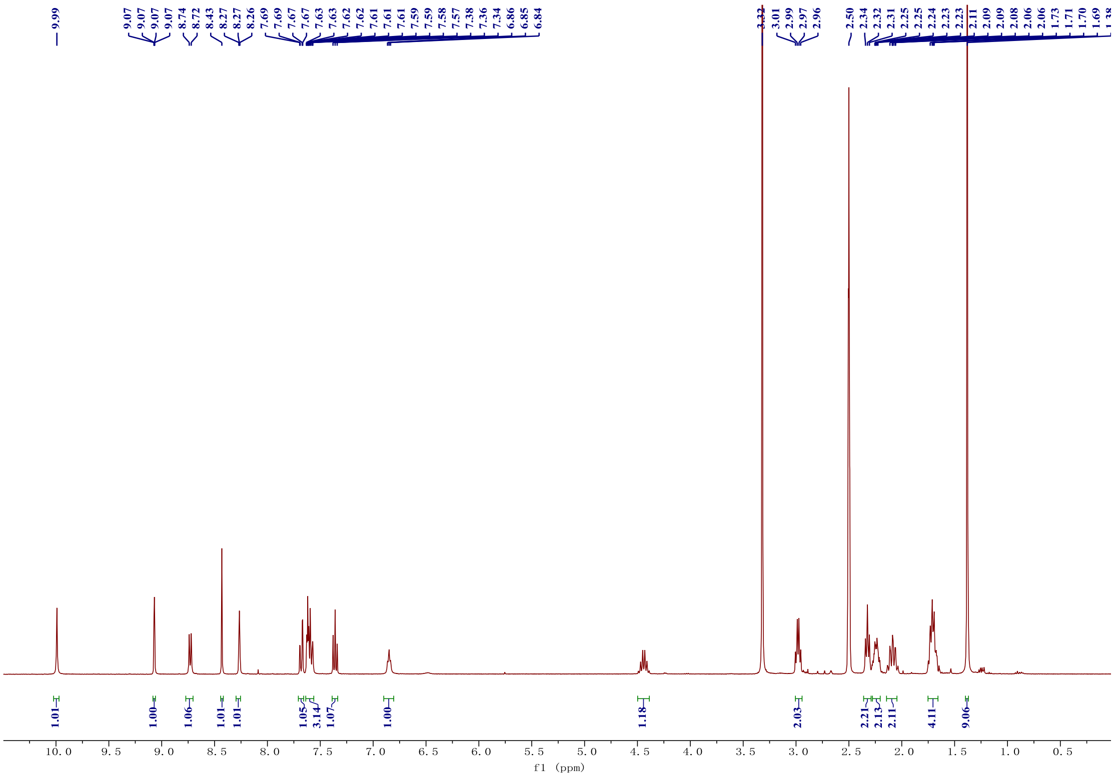

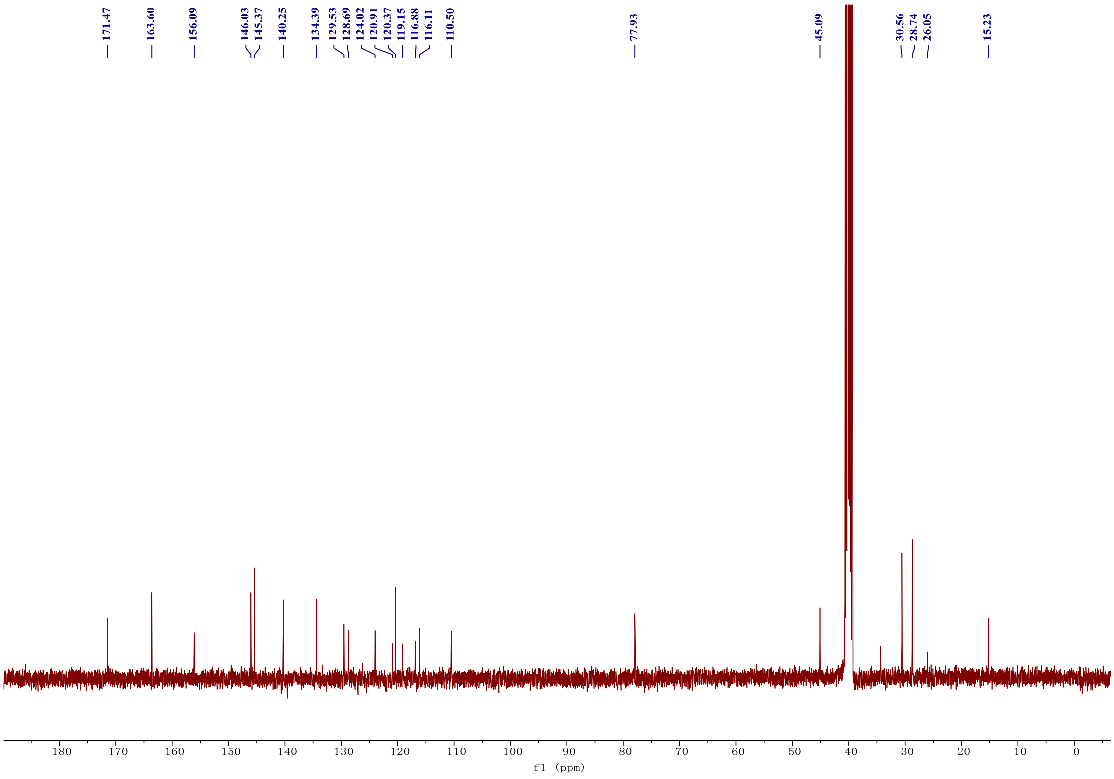

Figure S10. $^{1}H$ and $^{13}C$ NMR of compound **10** in DMSO-$d_6$.

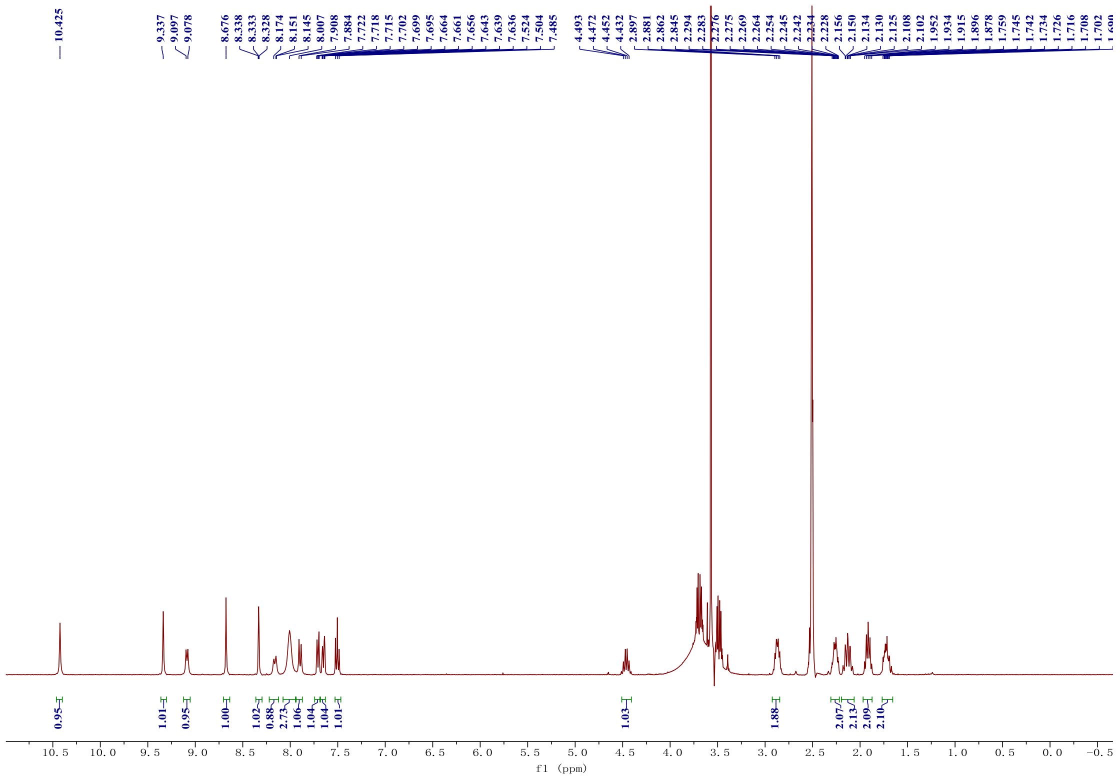

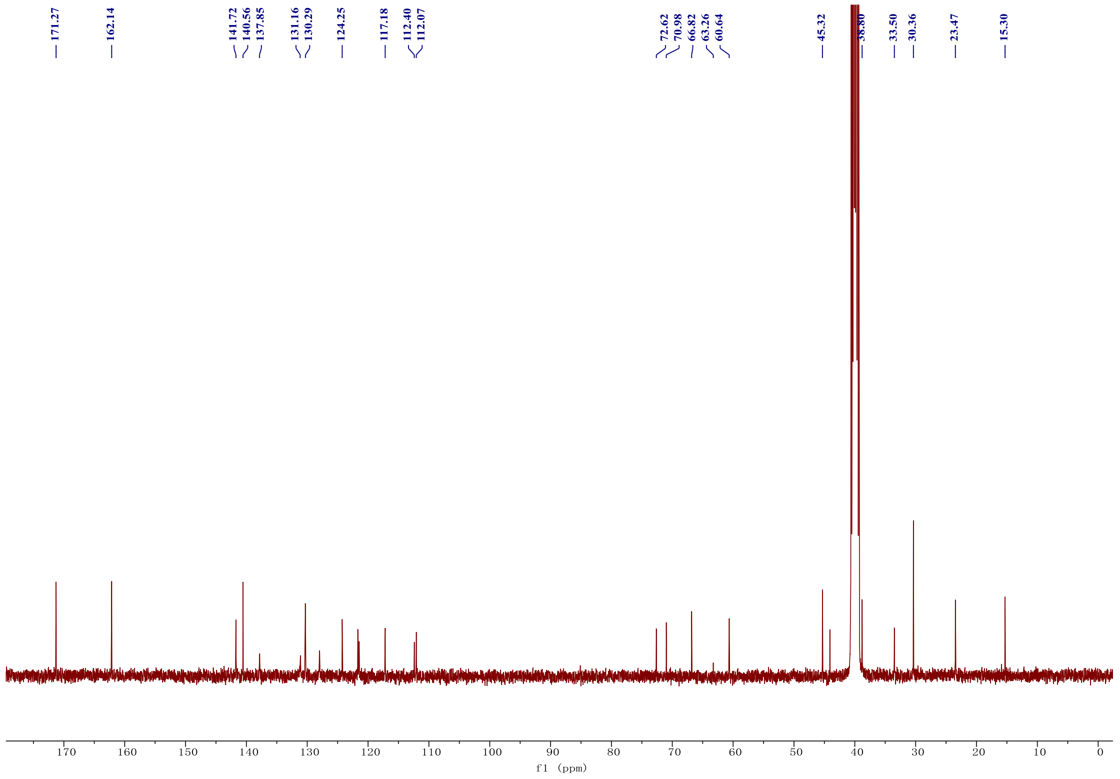

Figure S11. $^{1}H$ and $^{13}C$ NMR of compound **16** in DMSO-d$_6$.

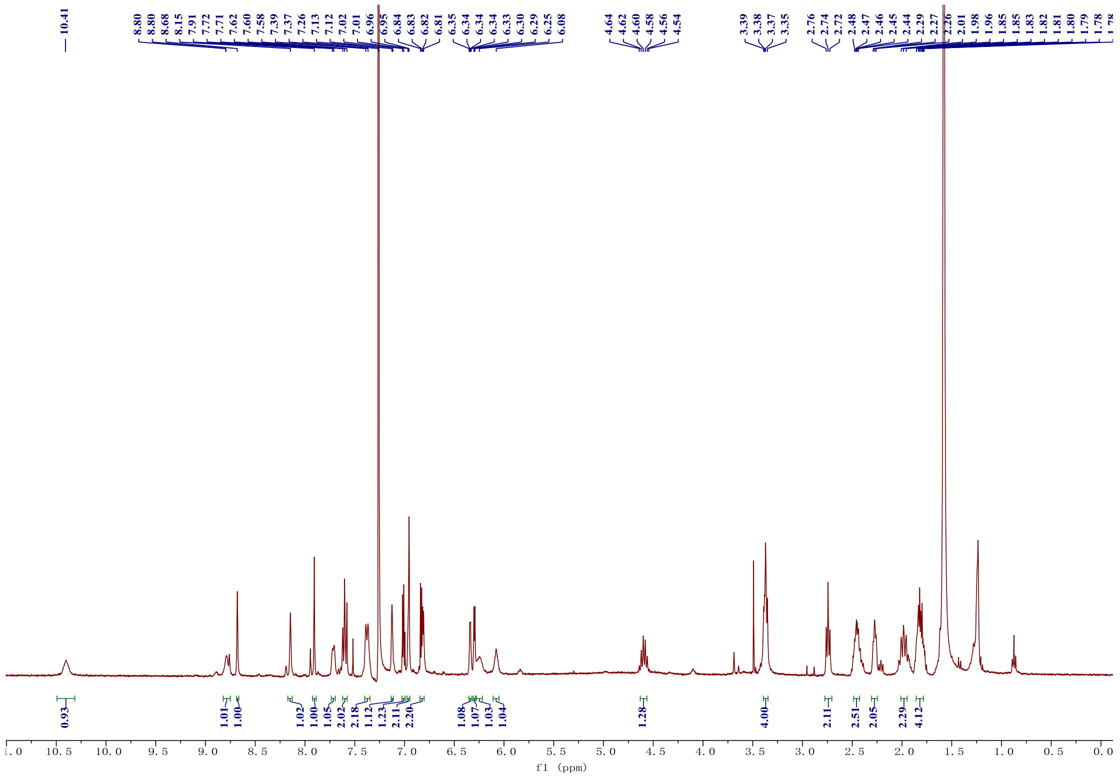

Figure S12. $^{1}$H NMR of **Tracer 2** in Chloroform-d.

1. Cribbs, A. P.; Kennedy, A.; Gregory, B.; Brennan, F. M., Simplified production and concentration of lentiviral vectors to achieve high transduction in primary human T cells. *BMC biotechnology* **2013,** *13* (1), 1-8.
2. Garnar-Wortzel, L.; Bishop, T. R.; Kitamura, S.; Milosevich, N.; Asiaban, J. N.; Zhang, X.; Zheng, Q.; Chen, E.; Ramos, A. R.; Ackerman, C. J.; Hampton, E. N.; Chatterjee, A. K.; Young, T. S.; Hull, M. V.; Sharpless, K. B.; Cravatt, B. F.; Wolan, D. W.; Erb, M. A., Chemical Inhibition of ENL/AF9 YEATS Domains in Acute Leukemia. *ACS Cent Sci* **2021,** *7* (5), 815-830.
3. Kim, H.; Kim, K.; Son, S.-H.; Choi, J. Y.; Lee, K.-H.; Kim, B.-T.; Byun, Y.; Choe, Y. S., 18F-Labeled BODIPY dye: A potential prosthetic group for brain hybrid PET/optical imaging agents. *ACS Chemical Neuroscience* **2018,** *10* (3), 1445-1451.
4. Robers, M. B.; Vasta, J. D.; Corona, C. R.; Ohana, R. F.; Hurst, R.; Jhala, M. A.; Comess, K. M.; Wood, K. V., Quantitative, real-time measurements of intracellular target engagement using energy transfer. In *Systems Chemical Biology*, Springer: 2019; pp 45-71.
5. Attwa, M. W.; Abdelhameed, A. S.; Alsaif, N. A.; Kadi, A. A.; AlRabiah, H., A validated LC-MS/MS analytical method for the quantification of pemigatinib: metabolic stability evaluation in human liver microsomes. *RSC Adv* **2022,** *12* (31), 20387-20394.
